# Skin but not gut microbial communities in Antarctic fur seals vary with social density

**DOI:** 10.1101/2025.03.19.644101

**Authors:** Petroula Botsidou, Michael Schloter, Öncü Maraci, Silvia Gschwendtner, Rebecca Nagel, Jaume Forcada, Joseph I. Hoffman

## Abstract

Comparative studies of microbial communities occupying different body sites in wild vertebrates are scarce, but they are crucial for advancing our understanding of the ecological and evolutionary factors shaping animal microbiomes. We therefore used a “natural experiment” comprising mother-offspring pairs from two adjacent Antarctic fur seal breeding colonies that differ in social density to investigate differences between skin and gut microbial communities in relation to host-specific and environmental factors. Using 16S rRNA amplicon sequencing, we uncovered a strong influence of colony on the diversity and composition of skin but not gut microbial communities. Specifically, we observed a suppressive effect of high social density on skin microbial alpha diversity as well as an overabundance of phyla associated with diseases and bite wounds in the high-density colony. Our findings suggest that skin microbial communities may be more sensitive to external factors, whereas gut communities are more tightly regulated by the host. Overall, this study highlights the importance of considering multiple body sites and their distinct microbial communities to develop a more comprehensive understanding of the factors shaping microbial diversity and composition in marine mammals.

## 1 Introduction

All vertebrates harbor complex and diverse microbial communities that play a crucial role in host physiology and survival (Coates et al., 2019). These microbial symbionts contribute to essential functions of the host including digestion, nutrient absorption, regulation of the immune system and the promotion of organ development and growth (Colston and Jackson, 2016; Dominguez-Bello et al., 2019; Zheng et al., 2020; Tan et al., 2022). Therefore, given the profound impact of the microbiota on host health and fitness, it is essential to understand the processes that shape host microbial communities.

The host-associated microbiota is specific to the body site it colonizes and can be influenced by a combination of host-specific and environmental factors (Coates et al., 2019). For example, the skin microbiota, which is continuously exposed to the external environment, is mainly influenced by external conditions while remaining distinct from the surrounding environmental microbes (Callewaert et al., 2020; Sehnal et al., 2021). Other factors such as age, sex, geographical location and maternal transfer are also known to shape skin microbial community composition and diversity (Ross et al., 2019; Sehnal et al., 2021; Howard et al., 2022). By contrast, diet is considered the primary determinant of gut microbial community composition (Colston and Jackson, 2016), although variation in the gut microbiota has been also linked to age (Pacheco-Sandoval et al., 2022), sex (Stoffel et al., 2020) and host genetics (Dabrowska and Witkiewicz, 2016).

Depending on the body site in question, host-related factors can also influence microbial community composition and diversity. At birth, human skin is first colonized by microbes from the mother’s birth canal and the surrounding environment (Coates et al., 2019; Luna, 2020). Skin microbial diversity then increases with age, mainly due to accumulated exposure to environmental microbes (Luna, 2020). However, this does not appear to be the case in wild mammals, with various studies suggesting that age is not associated with changes in skin microbial diversity (Chiarello et al., 2017; Lavrinienko et al., 2018; Grosser et al., 2019). By contrast, gut microbial communities of humans and wild mammals show little resemblance to their mother’s microbiota at birth and alpha diversity tends to increase with age, reflecting the dietary shift from milk to solid foods (Bäckhed et al., 2015; Baniel et al., 2022; Pacheco-Sandoval et al., 2022).

Among the various environmental factors influencing microbial communities, social density can have a strong effect on both the skin and gut microbiota. Higher density might promote higher microbial diversity by increasing the probability of microbial transmission, leading to a “social microbiome” (Li et al., 2016). However, crowded conditions can also induce physiological stress, which can negatively impact microbial abundance and function (Alverdy and Luo, 2017). For example, Antarctic fur seals breeding at high social density have significantly lower alpha diversity in their skin microbiota than conspecifics breeding at low social density (Grosser et al., 2019). Similarly, acute social stress has been linked to a decline in gut microbial diversity in Syrian hamsters (Partrick et al., 2018). Additionally, stress from overcrowding has been associated with changes in microbial community composition, often leading to inflammation and dysbiosis in both the skin and gut (Delaroque et al., 2021). Consequently, social density may be a key environmental driver of microbiome variation across different body sites.

More generally, the study of host-microbe-environment interactions is an emerging research field in wildlife biology that can offer valuable insights into animal ecology and health (Hoye and Fenton, 2018). However, most research focuses on the microbial communities of a single body site, with relatively few studies simultaneously examining multiple microbial communities within the same host. Comparative studies are therefore needed to produce a more comprehensive understanding of microbial diversity and to unravel the intricate interrelationships between hosts, their microbial symbionts and the environment (Martinez, 2018).

Pinnipeds, a charismatic group of marine mammals comprising the true seals, sea lions and the walrus, are ideally suited to investigating the effects of social density on the composition and diversity of skin and gut microbiomes. As semi-aquatic mammals, they inhabit both marine and terrestrial environments, exposing them to diverse microbial communities (Berta et al., 2018). Furthermore, many pinniped species breed in dense colonies, where close physical contact and social stress create optimal conditions for studying the effects of overcrowding on microbial diversity (Grosser et al., 2019).

The Antarctic fur seal (*Arctocephalus gazella*) provides a unique opportunity to investigate the effects of social density on the skin and gut microbiota of a wild, free-ranging marine mammal. This species breeds seasonally on sub-Antarctic islands, with around 96% of the global population concentrated around South Georgia in the southwest Atlantic (Forcada and Staniland, 2017). Pregnant females come ashore to give birth to their pups and rear them in breeding colonies that can vary in social density by an order of magnitude (Meise et al., 2016). On Bird Island, two breeding colonies–Freshwater beach (FWB) and Special study beach (SSB)–are located less than 200 meters apart and experience the same prevailing climatic conditions (Figure 1). Breeding females from both colonies share the same foraging grounds and do not differ significantly in body size or condition (Nagel et al., 2021). However, the density of breeding females is around four times higher at SSB relative to FWB (Meise et al. 2016) and, consequently, the density of pups is also markedly higher at SSB than FWB (Nagel et al., 2021). This is reflected by significantly higher levels of the stress hormone cortisol measured from the hair of mothers from SSB (Meise et al., 2016) and a suggestive yet weaker trend for cortisol measured from the saliva of mothers and pups to be higher at SSB (Nagel et al., 2022).

**Figure 1.**
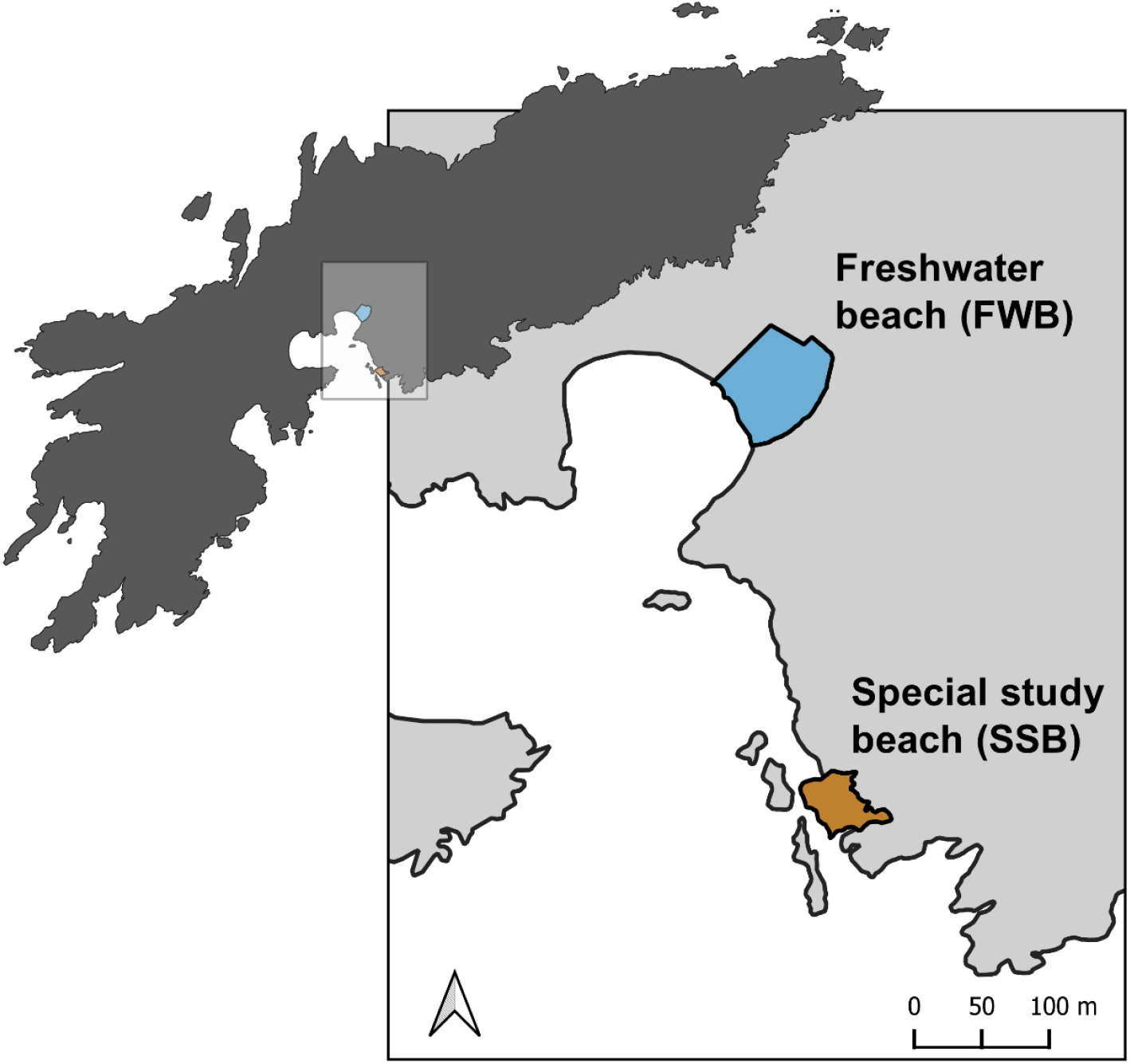
Map of the study area at Bird Island, South Georgia showing two adjacent Antarctic fur seal breeding colonies, Freshwater beach (FWB) and the Special study beach (SSB). The map was created using QGIS software version 3.32.2.

We took advantage of the “natural experiment” provided by these two breeding colonies to investigate the effects of host-specific and environmental factors on microbial community composition and diversity. Specifically, we used 16S rRNA sequencing to characterize the skin and gut microbial communities of 20 mother-pup pairs (total *n* = 40 individuals, 80 samples). We hypothesized that (i) mothers would carry similar skin microbial communities to their pups but exhibit greater gut microbial diversity, reflecting dietary differences between these two life-history stages; and (ii) microbial diversity in both skin and gut communities would be lower in animals sampled from SSB, consistent with the suppressive effects of high social density.

## 2 Methods

### 2.1 Study site and sample collection

Samples were collected from two adjacent Antarctic fur seal breeding colonies (Freshwater beach, FWB, and the Special study beach, SSB, Figure 1) at Bird Island, South Georgia (54°00’024.8 S, 38°03’04.1 W) during the austral summer of 2018–2019. Ten breeding females and their pups were captured from each colony. Methods used to restrain, and sample captured individuals followed protocols that have been established and refined as part of the long-term monitoring and survey program of the British Antarctic Survey (BAS). Each mother and her pup were captured concurrently around 2–3 days postpartum (in December). The mothers were captured with a noosing pole and held on a restraint board, while the pups were captured with a slip noose and restrained by hand. Biometric data (weight, length, span and girth) were gathered from the mothers and their pups; pups were also sexed.

Two microbial samples were collected from each individual. Skin microbial communities were sampled by rubbing a flocked swab (FLOQSwabs 552C, Copan Italia s.p.a.) across the animal’s cheek, underneath the eye and behind the snout, while gut microbial communities were sampled by swabbing inside the anus. The resulting microbial samples (*n* = 80) were individually stored in sterile transport tubes at −20°C during the field season and subsequently at −80°C in the laboratory. Additionally, environmental controls (i.e. swabs from the soil and air, as well as from the gloves and skin of the animal handlers) were collected (*n* = 6) with the aim of removing potentially contaminating sequences from our samples. A full list of the samples that were collected and analyzed in this study are available in Supplementary table S1.

### 2.2 DNA extraction and 16S rRNA gene sequencing

Microbial DNA was extracted from all 86 swabs using a NucleoSpin DNA Forensic Isolation Kit (Macherey-Nagel, Düren, Germany). We followed the manufacturer’s recommended protocol with the exception that we eluted the DNA in 45 μL of FOE buffer that was preheated to 70°C. The library preparation and 16s rRNA amplicon sequencing were conducted by Biomarker Technologies (BMK) GmbH. A ∼247-bp region of the V3–V4 region of the 16S rRNA gene was amplified using the primers 338F (5’ – ACTCCTACGGGAGGCAGCA – 3’) and 806R (5’ – GGACTACHVGGGTWTCTAAT – 3’). The polymerase chain reaction (PCR) mixture contained 2.5–4 ng of DNA, 0.3 μl of each primer (diluted to 10 μM), 5 μL of KOD FX Neo Buffer (Beijing Biolink Biotechnology Co., Ltd), 2 μL dNTPs, 0.2 μL KOD FX Neo (Beijing Biolink Biotechnology Co., Ltd) and ddH_2_0 up to a total volume of 10 μL. The PCR amplification program comprised an initial denaturation step of 95°C for 5 min followed by 25 cycles of denaturation at 95°C for 30 s, annealing at 50°C for 30 s, extension at 72°C for 40 s and a final extension step at 72°C for 7 min. The PCR products were then purified using VAHTSTM DNA Clean Beads (Nanjing Vazyme Biotechnology Co., Ltd.) and amplified in a 20 μL indexing PCR consisting of 5 μL of purified PCR product, 2.5 μL of each indexing primer (Mpp1-n1, Mpp1-n2, diluted to 10 μM; Suzhou SYNBIO Technology Co., Ltd.) and 10 μL Q5 High Fidelity 2x Master Mix. The indexing PCR amplification program comprised an initial denaturation of 98°C for 30 s followed by 10 cycles of 98°C for 10 s, 65°C for 30 s, 72°C for 30 s and a final extension step of 72°C for 5 min. PCR products were run on a 1.8% agarose gel and quantified with the ImageJ software. 150 ng of each sample was then pooled, purified with E.Z.N.A.® Cycle Pure Kit (Omega; Beijing Hongyue Innovation Technology Co., Ltd.) and size-selected by gel electrophoresis with the Monarch DNA Gel Extraction Kit (Beijing Hongyue Innovation Technology Co., Ltd.). The library was then paired-end sequenced on an Illumina Novaseq 6000 sequencing platform.

### 2.3 Sequence data processing

Paired-end sequences (247bp) were analyzed on the Galaxy web platform (www.galaxy.org; Afgan et al., 2016). First, adapters were trimmed using Cutadapt (Martin, 2011) and only reads with a minimum read length of 50 were retained. Quality control was performed using FastQC (Andrews, 2010). All of the samples contained adequate numbers (>10,000) of high-quality reads, and we therefore retained them all for subsequent analyses. DADA2 (Galaxy version 1.28; (Callahan et al., 2016) was used for further filtering and processing the raw sequences into amplicon sequence variants (ASVs) after trimming 20 bp from the 5’ end of each read and setting the expected error to three for the forward reads and four for the reverse reads. Taxonomy was assigned to the ASVs using the SILVA database version 138.1 (Quast et al., 2013) and the phyla names were then corrected according to Oren and Garrity, 2021. Using the decontam package version 1.18.0 (Davis et al., 2017) in R version 4.3.1 (R Core Team, 2022) with the “prevalence” method setting the probability threshold to 0.1, a total of 421 potential contaminant ASVs were removed for the skin and 699 for the gut. In addition, ASVs classified as mitochondrial or chloroplastic, which could not be identified at the phylum level, or which were present in only one sample (singletons) were discarded. Microbial community composition and the core microbiota were characterized for both the skin and gut samples, with the latter defined to include only ASVs present in at least 90% of the samples. This analysis was performed at the phylum level based on non-normalized ASVs. The resulting ASV tables were subsequently analysed in R version 4.3.1 (R Core Team, 2022).

### 2.4 Statistical analyses

We calculated microbial alpha diversity using the Shannon diversity index, which considers both the richness and evenness of the taxa present (Allaby, 2014). Shannon diversity was computed using the phyloseq R package version 1.42.0. To control for variation in sequencing depth among the samples, the reads were rarefied to the lowest read depth, following observations of rarefaction curves (Supplementary figure S1). However, the resulting alpha diversity metrics were strongly positively correlated with those based on the non-rarefied dataset (Spearman’s rank correlation: *r* = 0.99, *p* < 0.001). Consequently, in order to maximise the available data analysed and to avoid excluding rare ASVs, we used the non-rarefied dataset for subsequent analysis.

We aimed to investigate the effects of body site (skin versus gut), colony (FWB versus SSB) and age class (mother versus pup) on alpha diversity. First, we compared the Shannon diversity index between skin and gut samples, finding them to be significantly different (Kruska-Wallis: *χ*^*2*^_(1)_ *=* 47.4, *p* < 0.001, Supplementary figure S2). We then fitted two linear mixed effect models (LMMs) with the lmerTest R package version 3.1-3 (Kuznetsova et al., 2017); we ran separate models for skin and gut microbial communities. Each model included Shannon index as the response variable and colony and age class as fixed effects. Mother-pup pair ID was included as a random effect to control for the spatial proximity of mothers and their offspring. To explore the effects of sex on alpha diversity, we also ran linear models (LM) only for pups, which included both female and male observations. For all models, the significance of the fixed effects was assessed at *a* = 0.05 by running Type II and Type I analysis of variance (for the LMs and LMMs respectively), as well as by evaluating whether the 95% confidence intervals (95% CIs) of the effect size estimates overlapped zero. The significance of the random effects was assessed using the ranova function of the lmerTest R package. The variance explained by the fixed effects (*R*^2^_marginal_) and by both the fixed and random effects (*R*^2^_conditional_) was calculated using the r.squaredGLMM function of the MuMIn package version 1.47.5 (Barton, 2023). The model residuals were inspected for normality and homogeneity of variance using the performance package version 0.12.0 (Lüdecke et al., 2021).

Before testing for differences in microbial composition (beta diversity) among the samples, we applied cumulative sum scaling (CSS) normalization our filtered dataset using the metagenomeseq package version 1.30.0 (Paulson et al., 2013). We then calculated pairwise microbial dissimilarity matrices based on Bray-Curtis dissimilarities and visualized differences among the groups using non-metric multidimensional scaling (nMDS) plots. To statistically assess whether any differences in microbial composition could be related to colony, age class or mother-pup pair, we implemented permutational multivariate analyses of variance (PERMANOVA; Anderson, 2017) with 1,000 permutations using the adonis function of the vegan R package version 2.6-4 (Oksanen et al., 2017). To evaluate whether significant results from the PERMANOVA arose due to variation in dispersion within groups instead of from differences in the mean values, we performed homogeneity of group dispersion tests using betadisper in the vegan R package (Oksanen et al., 2017).

Finally, we identified differentially abundant ASVs between colonies and age classes by analysing the filtered and non-normalized datasets using the corncob package version 0.4.1 (Martin et al., 2020). This software uses beta-binomial regressions to model the relative abundances of all the taxa while taking into account differential variability. Significance testing was performed using likelihood ratio tests and the *p*-values were obtained after controlling for the Type I error using Bonferroni correction (*p* < 0.05).

## 3 Results

We characterized the skin and gut microbial communities of 20 Antarctic fur seal mother-pup pairs from two adjacent breeding colonies of contrasting social density (FWB = low density, SSB = high density, Figure 1). After stringent quality filtering, we retained a total of 4,546,491 reads for the skin (mean = 113,662.3 *±* 19,627 s.d. reads per sample) and 4,923,677 reads for the gut (mean = 123,091.9 ± 11,273.4 s.d. reads per sample) (Supplementary tables S2-S4). These were processed and clustered into 6,946 ASVs for the skin (mean *=* 1,623.7 ± 345 s.d. ASVs per sample) and 3,719 amplicon sequence variants (ASVs) for the gut (mean *=* 685.8 ± 268.7 s.d. ASVs per sample) as shown in Supplementary table S4.

### 3.1 Antarctic fur seal microbiota composition

We identified 30 distinct microbial phyla for the skin and 25 for the gut (Figure 2, Supplementary table S5). Although both body sites shared the same five most abundant phyla, their relative abundances differed (Pseudomonadota: skin = 30.1%, gut = 18.3%; Bacillota: skin = 26%, gut = 36.2%; Actinomycetota: skin = 17.4%, gut = 8.2%, Bacteroidota: skin = 14.6%, gut = 16%; Fusobacteriota: skin = 8%, gut = 15.6%). Focusing on those taxa present in at least 90% of the samples from each body site, we identified 136 core ASVs in the skin and 64 in the gut (Supplementary tables S6 and S7). Four phyla (Bacillota, Pseudomonadota, Fusobacteriota and Actinomycetota) dominated the core microbiota of both the skin and the gut, with Pseudomonadota being the most abundant phylum in the skin core microbiota (33%) and Bacillota being the most abundant phylum in the gut core microbiota (42%).

**Figure 2.**
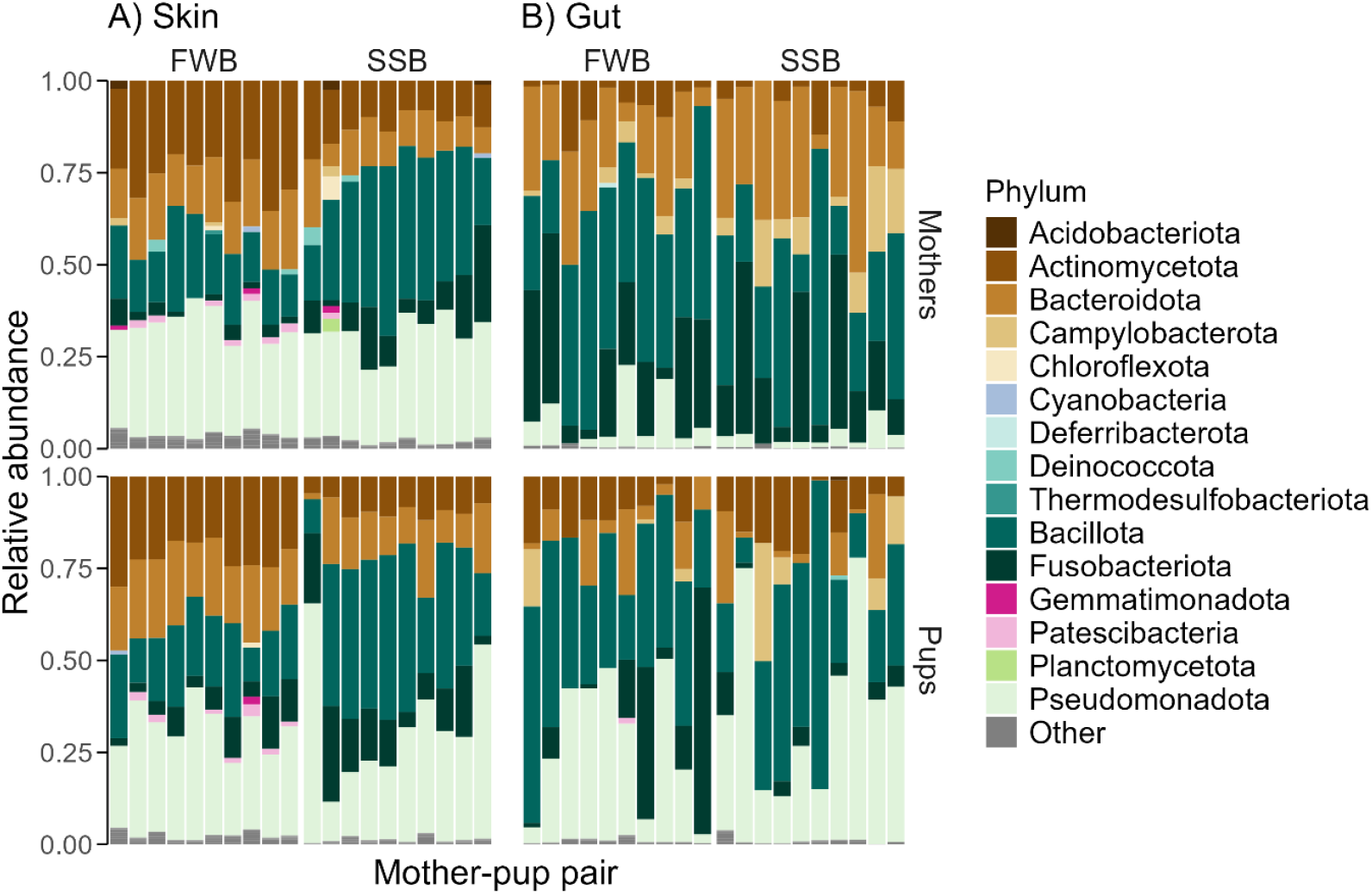
The relative abundances of common microbial phyla in 80 Antarctic fur seal samples grouped by colony (Freshwater beach, FWB, versus Special study beach, SSB) and age class (mothers versus pups). Only those phyla with relative abundances above 1% are shown, with the rest categorized as “Other”. The identities of mother-pup pairs are indicated by their placement in the figure, with each mother being plotted directly above her pup.

### 3.2 Patterns of alpha and beta diversity

Skin microbial alpha diversity, quantified using the Shannon index, was significantly higher than gut alpha diversity (Kruska-Wallis: *χ*^*2*^_(1)_ *=* 47.4, *p* < 0.001) and there was little overlap in the distributions of Shannon index values between the two body sites (Supplementary figure S2). For the skin, microbial alpha diversity was significantly lower in the high-density colony (SSB; LMM_Colony = SSB_: *β* = −0.74, *χ*^*2*^ *=* 17.51, *p <* 0.001), but did not differ significantly between mothers and pups (LMM_Age class = Pup_: *β* = −0.24, *χ*^*2*^ *=* 3.08, *p =* 0.08, Figure 3A, Supplementary table S8), with the fixed effects explaining almost 40% of the variation (*R*^*2*^_*marginal*_ = 0.38, *R*^*2*^_*conditional*_ *=* 0.52). By contrast, gut alpha diversity did not differ significantly between the colonies (LMM_Colony = SSB_: *β* = −0.07, *χ*^*2*^ = 0.09, *p* = 0.76) or between mothers and pups (LMM_Age class = Pup_: *β* = 0.06, *χ*^*2*^ = 0.07, *p =* 0.79; Figure 3C, Supplementary table S8), with the fixed effects explaining only a small proportion of the total variation (*R*^2^_marginal_ *=* 0.004, *R*^2^_conditional_ = 0.01). Lastly, no differences in alpha diversity were found between male and female pups for either the skin (LM_Sex = Male_: *β* = −0.51, *χ*^*2*^ = 3.36, *p* = 0.08) or gut (LM_Sex = Male_: *β* = −0.02, *χ*^*2*^ = 1e-03, *p* = 0.98; Supplementary table S9).

**Figure 3.**
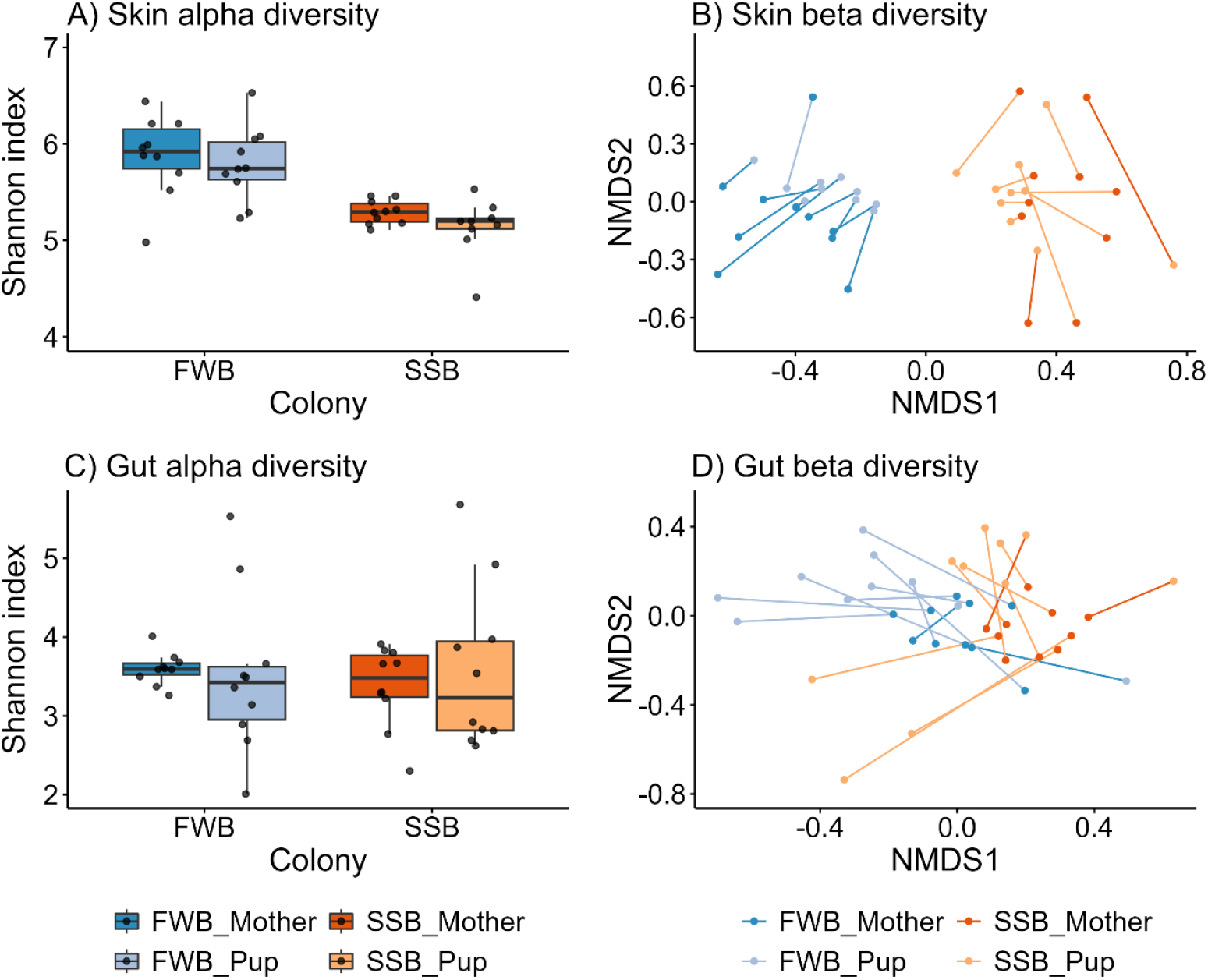
Alpha (A, C) and beta (B, D) diversity shown separately for microbial communities of the skin (A, B) and gut (C, D). Alpha diversity was calculated using the Shannon index on non-rarefied ASVs and plotted by breeding colony (Freshwater beach, FWB, versus Special study Beach, SSB) and age class (mothers versus pups). Beta diversity differences in relation to breeding colony and age class were visualized using non-metric multidimensional scaling (nMDS) plots of Bray-Curtis dissimilarities. The lines connect individual mothers to their pups. For clarity, we removed a single outlier (pup H11) from panel (A) (Supplementary figure S3) although this individual and its mother were retained in all of the analyses. The horizontal line within each box plot represents the median, while the lower and upper boundaries of the boxes indicate the 25^th^ and 75^th^ percentiles, respectively, encompassing the interquartile range (IQR). Whiskers extend from the boxes to a maximum of 1.5 times the IQR.

Visualizing microbial composition with nMDS plots based on Bray-Curtis dissimilarities, we observed two distinct clusters corresponding to FWB and SSB for the skin, while clustering was less defined for the gut (Figure 3B, Figure 3D). This was reflected in the results of PERMANOVAs, which revealed an almost three times higher proportion of the variance explained by colony for the skin (PERMANOVA_Colony_: *R*^2^ = 0.23, *p* = 0.001) relative to the gut (PERMANOVA_Colony_: *R*^2^ = 0.08, *p* = 0.001; Supplementary table S10). Significant differences in microbial composition between the age classes were found for both body sites (PERMANOVA_Skin-Age class_: *R*^*2*^ = 0.04, *p* = 0.009; PERMANOVA_Gut-Age class_: *R*^*2*^ = 0.09, *p =* 0.001) although this pattern was weaker for the skin (Supplementary table S10). Additionally, microbial communities showed significantly greater variability among mothers for the skin (betadisper_Skin-Age class_: *F* = 11.13, *p* = 0.002) and among pups for the gut (betadisper_Gut-Age class_: *F* = 18.6, *p* < 0.001; Figure 3B, Figure 3D).

As previously found by Grosser et al. (2019), mothers and their pups also exhibited significantly more similar skin microbial communities than expected by chance (PERMANOVA_Skin-PairID_: *R*^*2*^ = 0.45, *p* = 0.001). However, this pattern was not present in the gut microbial communities (PERMANOVA_Gut-PairID_: *R*^*2*^ = 0.42, *p* = 0.11). Additionally, no significant differences in microbial composition were found between male and female pups either for the skin or the gut (PERMANOVA_Skin-Sex_: *R*^*2*^ = 0.05, *p* = 0.24; PERMANOVA_Gut-Sex_: *R*^*2*^ = 0.06, *p* = 0.27; Supplementary table S10).

### 3.3 Differentially abundant taxa

Using beta binomial regression models, we identified differentially abundant ASVs between the two colonies and age classes. A greater number of differentially abundant ASVs was detected for the skin compared to the gut in relation to both colony and age class. For colony comparisons, most of the differentially abundant ASVs were more abundant in the low-density colony (FWB) and corresponded to the phyla Patescibacteria, Gemmatimonadota, Bacteroidota and Actinomycetota. By contrast, only two ASVs were more abundant in the high-density colony (SSB), belonging to the Fusobacteriota and Bacillota (Figure 4A). Regarding age class, no differentially abundant ASVs were identified for pups, while the taxa that were more abundant in mothers included phyla typically present in diverse terrestrial and marine habitats, as well as the Campylobacterota (Figure 4B).

**Figure 4.**
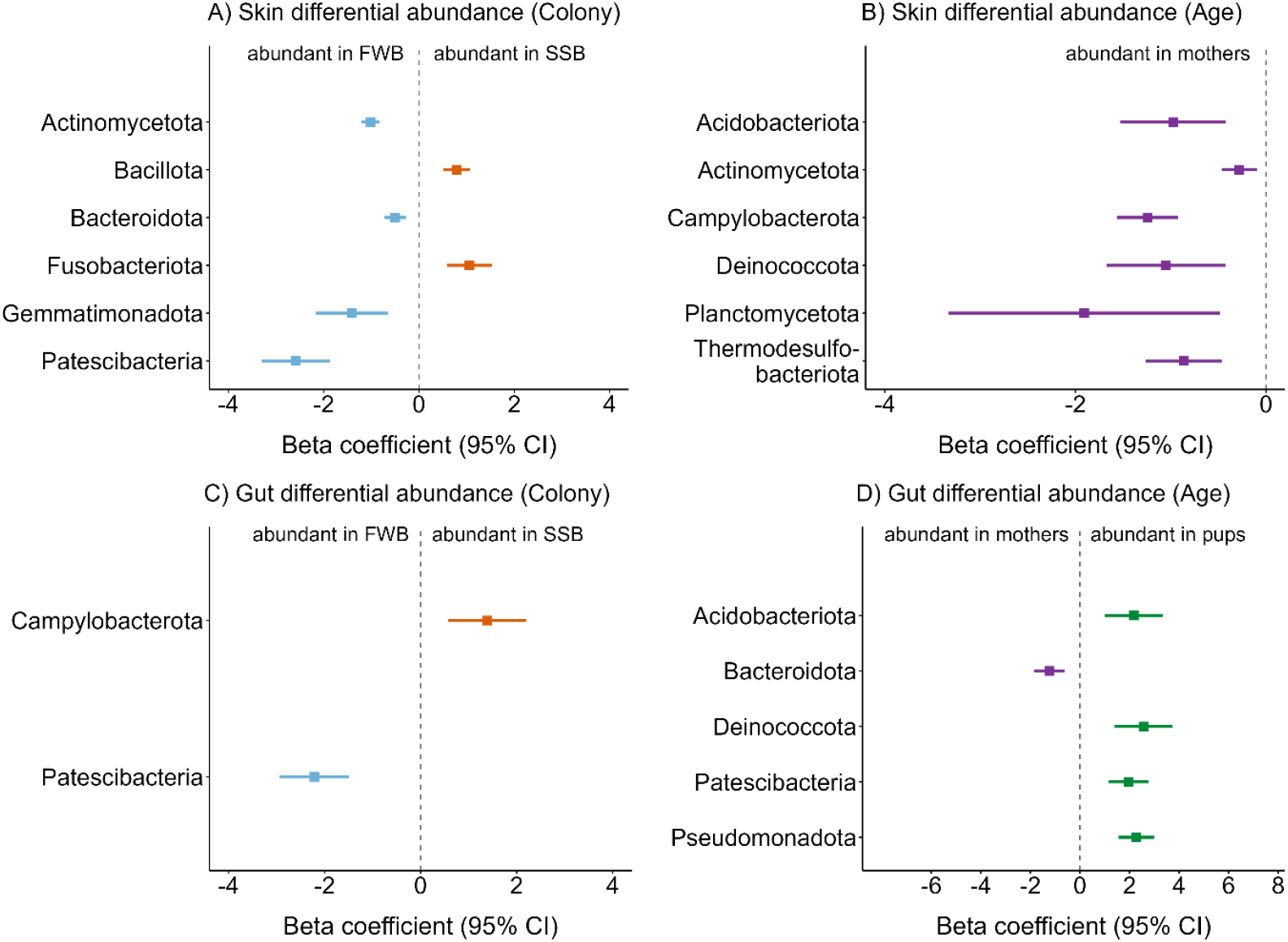
Differentially abundant ASVs in relation to breeding colony (A, C) and age class (B, D) for skin (A, B) and gut (C, D) microbial communities. The phylum-level taxonomy is presented for each corresponding ASV. Differential abundance analysis was performed by constructing beta binomial regression models as described in the Methods. Estimates are plotted with ±95% confidence intervals (CI).

For the gut, only two ASVs were differentially abundant between the two colonies: An ASV that was more abundant in the high-density SSB belonged to the Camplylobacterota, a phylum associated with bite wounds in pinnipeds (Foster et al., 2020), while an ASV that was more abundant in the low-density FWB belonged to the Patescibacteria (Figure 4C). Most of the differentially abundant ASVs were detected in relation to age class for the gut. Only one ASV belonging to the Bacteroidota was significantly more abundant in mothers, with the rest of the pup-associated ASVs corresponding to phyla commonly found in the soil, freshwater and marine environments, as well as the Pseudomonadota, a taxon that is detected in human infants (Yang et al., 2024, Figure 4D).

## 4 Discussion

Comparative studies of microbial communities across different body sites in wild vertebrates are crucial for understanding the ecology and evolution of host associated microbiota. However, most studies focus on single body sites, overlooking the potential for complex interactions between ecological and evolutionary factors and the microbial communities of different tissues within a single host. We therefore used skin and gut swabs from the same individuals to investigate how host-specific and environmental factors shape the skin and gut microbial communities of Antarctic fur seals. We found that, despite sharing core phyla, the skin and gut microbiota exhibited distinct responses to environmental and host-related factors. Notably, colony strongly influenced the alpha and beta diversity of skin but not gut microbial communities. By contrast, age class had a weak effect on skin and gut beta diversity, whereas sex did not significantly affect either alpha or beta diversity in the skin or gut microbiota. Our findings emphasize the differential impacts of host-specific and environmental factors on distinct microbial ecosystems within an individual.

### 4.1 Composition of the skin and gut microbiota

We found that the skin and gut microbial communities of Antarctic fur seals were distinct, despite sharing five highly abundant phyla (Bacillota, Pseudomonadota, Bacteroidota, Fusobacteriota and Actinomycetota). More specifically, the phylum Pseudomonadota dominated the skin microbial communities. This was expected as this taxon is generally detected at high abundance on the skin of cetaceans and fur seals, as well as in seawater (Grosser et al., 2019; Van Cise et al., 2020). By contrast, and in line with other studies of Antarctic pinnipeds, including other species of fur seals (Nelson et al., 2013; Medeiros et al., 2016), the gut microbiota showed a high abundance of Bacillota. This taxon, a well-known gut colonizer, is thought to aid the host in efficiently extracting energy from food, which promotes fat deposition, an essential process for pinnipeds that depend heavily on fat storage for energy intake and thermoregulation (Pacheco-Sandoval et al., 2019).

### 4.2 The effects of breeding colony on alpha and beta diversity

The environment in which a host lives, including both the social and physical environment, can have a major impact on host microbial communities (Tasnim et al., 2017; Callewaert et al., 2020). To investigate the effects of host and environmental factors on the Antarctic fur seal microbiota, we therefore used a unique natural experiment comprising two adjacent breeding colonies that differ in social density but which experience ostensibly the same environmental conditions due to their close physical proximity. Focusing initially on skin microbial communities, we found marked differences between the two colonies, with alpha diversity being significantly lower in the high-density colony. This corroborates the results of a previous study of mother-pup pairs from the same two colonies that used similar approaches but only investigated the skin microbiota (Grosser et al., 2019). As these two studies were conducted four years apart and there was no overlap between the individuals sampled at the two timepoints, we conclude that these patterns are reproducible and hence show temporal stability. This finding is important in its own right as replication studies in the fields of ecology and evolution are uncommon (Kelly et al 2006; Nakagawa & Parker, 2015), reflecting the general perception that research in these fields can be difficult to replicate, partly due to the challenges of controlling for confounding factors such as environmental variation in natural settings.

Clear and repeatable differences in skin microbial alpha diversity between the high- and low-density colony are consistent with a suppressive effect of social density on microbial diversity (Grosser et al., 2019). In line with this, cortisol levels have been shown to be elevated in mothers and pups from SSB (Meise et al., 2016; Nagel et al., 2022) reflecting a potentially more stressful and competitive environment. A negative association between elevated social stress and alpha diversity has been also reported in captive hamsters (Partrick et al., 2018), while other studies have shown that high stress levels, indicated by increased glucocorticoid concentrations, have a suppressive effect on alpha diversity in humans and red squirrels (Stothart et al., 2016; Keskitalo et al., 2021). Therefore, our results lend further support to the argument that overcrowding increases social stress and reduces skin microbial alpha diversity.

By contrast, gut alpha diversity did not differ significantly between FWB and SSB. Even though studies of the impacts of social density and stress on mammalian gut microbiomes are scarce, a similar pattern was observed in a recent study of laboratory mice in which social overcrowding elevated corticosterone levels without affecting gut alpha diversity (Delaroque et al., 2021). Similarly, studies of great apes (Vlčková et al., 2018; Hickmott et al., 2022) did not find any significant effects of stress on gut microbial diversity, suggesting that these species may buffer stress through social interactions and behavioral flexibility (Hickmott et al., 2022). Overall, the lack of an obvious relationship between proxy measures of social stress and gut alpha diversity in Antarctic fur seals and other species may be a reflection of the host’s ability to regulate its microbiota, maintaining homeostasis and protecting against harmful colonizers (Foster et al., 2017). In particular, gut epithelial cells play a central role by secreting antimicrobial peptides and limiting oxygen availability, thereby fostering a microbiota that supports host health and nutrition (Dabrowska and Witkiewicz, 2016).

To further investigate community-level differences, we analyzed the beta diversity of the two breeding colonies across both body sites. Consistent with our results for alpha diversity, we found pronounced differences in microbiota composition between the colonies for the skin, while these differences were considerably weaker for the gut. This alignment between alpha and beta diversity is not surprising as the composition and diversity of microbial communities are often closely linked (Grosser et al., 2019). This linkage suggests that similar regulating factors may influence both aspects of microbial community structure.

### 4.3 Microbial community differences between mothers and pups

Our dataset also enabled us to test for differences in microbial communities between mothers and pups. We found no significant differences in alpha diversity between the two age classes for either the skin or the gut. Similarly, while mothers and pups exhibited significant differences in microbial community composition, these patterns were relatively weak, particularly for the skin. These findings are consistent with a previous study of Antarctic fur seals that also observed minimal differences in skin microbial diversity and composition between mothers and pups from the same colonies (Grosser et al., 2019). However, they contrast with findings from studies of other pinnipeds, such as harbor seals and southern elephant seals, where pronounced differences in the gut microbiota of adult females and pups were observed (Nelson et al., 2013; Pacheco-Sandoval et al., 2022). The similarities between Antarctic fur seal mothers and newborn pups may result from the vertical transmission of microbes at birth, during nursing and via close physical proximity, which also exposes them to similar environmental microbes (Coates et al., 2019). By contrast, Pacheco-Sandoval et al. (2022) observed pronounced differences in the gut microbiota of harbor seal mothers and pups, likely due to diet-driven changes as the pups were transitioning from milk to solid foods. To better understand the colonization and development of microbial communities during early life-stages, longitudinal studies of Antarctic fur seals pups from birth to weaning will be needed.

Additionally, we found that mother-pup pairs shared more similar microbial communities than expected by chance for the skin (as previously reported by Grosser et al., 2019) but not for the gut. As argued above, these similarities in skin microbial communities are to be expected given the close proximity of mothers and their newborn pups during activities such as suckling and resting, which can facilitate the transmission of skin microbes (Coates et al., 2019). By contrast, the development of the gut microbiota may be more strongly influenced by factors such as dietary shifts, including the transition from milk to solid food, as well as by the maturation of the immune system (Coates et al., 2019; Reese et al., 2021).

### 4.4 Taxon-specific differences between colonies and age classes

To gain deeper insights into the patterns described above, we conducted differential abundance analyses. Focusing on the effects of social density, we found that several taxa typically associated with pathogenesis were enriched in both the skin and gut microbiota of individuals from the high-density colony (SSB). Notably, Bacillota were more abundant in the skin microbiota of animals from SSB, a taxon previously linked to skin wounds and disease progression in stranded porpoises (Li et al., 2022). Overcrowding and social stress are likely contributing factors, as they are known to reduce microbial alpha diversity and induce shifts in microbial abundance that can lead to dysbiosis and the colonization of pathogenic taxa (Alverdy and Luo, 2017; Delaroque et al., 2021).

Interestingly, despite the absence of significant differences in gut alpha diversity between the colonies and only a weak influence of breeding colony on gut beta diversity, we found a significantly higher abundance of Campylobacterota in the gut microbiota of individuals from the high-density colony (SSB). This phylum has been linked to gastrointestinal diseases in humans and to pathogenesis associated with bite wounds in pinnipeds (Foster et al., 2020; Watkins et al., 2022). This observation lends further support to the argument that, while animals from both colonies maintain diverse microbiota, individuals from the high-density colony appear to experience an enrichment of pathogenic taxa. To better understand the effects of social stress on the gut microbiome, future studies should incorporate measurements of stress hormones such as cortisol to explore the relationship between stress and the gut microbiota at the individual level. Additionally, metagenomic approaches could be used to investigate the functional characteristics of differentially abundant microbes and their roles in host health and disease.

Above and beyond these patterns, we also identified several differentially abundant taxa at the low-density colony (FWB) belonging to phyla commonly found in soil, freshwater and marine habitats (i.e. Patescibacteria, Actinomycetota, Gemmatimonadota, Haro-Moreno et al., 2023; Mujakić et al., 2022; Anandan et al., 2016). The presence of a small stream at FWB likely facilitates the acquisition of these environmental microbes, which appear to be an important source of variability in microbial diversity and composition at the low-density colony. This finding aligns with a previous study of harbor seals, where populations near freshwater sources were found to exhibit higher alpha diversity (Pacheco-Sandoval et al., 2019). It emphasizes the importance of environmental sources of variation, particularly in the absence of external stressors such as overcrowding, and further supports the idea that host habitat plays a key role in shaping microbial community composition in marine mammals, alongside diet and phylogeny (Bik et al., 2016).

Finally, regarding age class, we found that the gut microbiota of pups was dominated by the phyla Pseudomonadota, while gut microbiota of mothers was dominated by Bacteroidota. This pattern has been previously described in human mothers and their infants (Suárez-Martínez et al., 2023; Yang et al., 2024). It reflects the natural progression of gut colonization during development, with facultative anaerobes like Pseudomonadota thriving in the oxygen-rich environment of the newborn gut. As individuals transition from milk to solid foods during weaning, the gut microbiota becomes dominated by phyla like Bacteroidota, which are able to break down complex polysaccharides (Suárez-Martínez et al., 2023). We also found that the gut microbiota of pups was enriched for environmental phyla such as Patescibacteria, Deinococcota, and Acidobacteriota, suggesting that many early colonizers originate from the environment where the individuals are born (Schmidt et al., 2019; Suárez-Martínez et al., 2023; Yang et al., 2024).

## 5 Conclusion

In conclusion, our study explored the interplay between host-specific and environmental factors in shaping the skin and gut microbial communities of Antarctic fur seals. We observed strong colony-level differences in skin microbial communities as well as clear differentiation among mother-pup pairs. However, these differences were less pronounced or absent in the gut microbiota of the same individuals. We also identified differentially abundant taxa between the colonies, including both potential pathogens and taxa commonly acquired from the environment. This study provides insights into how ecological factors, such as social density, shape the composition and diversity of microbial communities occupying different body sites in a wild vertebrate. We recommend that future research should build on these findings by characterizing the functional consequences of host-microbe interactions in Antarctic fur seals, as well as by including additional factors such as host physiology and genetics. This should provide more comprehensive insights into how microbes affect the health and fitness of Antarctic fur seals in heterogenous environments.

## Data availability statement

The 16S raw reads that have generated for this study are deposited at the European Nucleotide Archive (ENA) under project accession number PRJEB86671. The scripts and metadata for reproducing the analyses can be accessed via the GitHub repository: https://github.com/ecopetridish/AFS_Skin_And_Gut_Microbiome

## Ethics statement

The Antarctic fur seal samples used in this project were collected and retained under permits issued by the UK’s Department for Environment, Food and Rural Affairs (Animal Health Act, import license number ITIMP18.1397). All fieldwork procedures were approved by the British Antarctic Survey Ethics Committee (AWERB applications 2018/1050 and 2019/1058), which fully complies with Home Office regulations and guidelines. The work was also carried out under the Scientific Research Permit for the British Antarctic Survey field activities on South Georgia, issued by the Government of South Georgia and the South Sandwich Islands (Wildlife and Protected Areas Ordinance (2011), RAP permit numbers 2018/ 024 and 2019/032). In addition, all aspects of the work covered by this project were in full adherence with the Convention on International Trade in Endangered Species of Wild Fauna and Flora (import nos 578938/01-15 and 590196/01-18).

## Author’s contributions

PB: Conceptualization, Data curation, Formal analysis, Investigation, Methodology, Software, Validation, Visualization, Writing – original draft, Writing – review & editing. MS: Conceptualization, Funding acquisition, Resources, Supervision, Validation, Writing – review & editing. ÖM: Investigation, Validation, Writing – review & editing. SG: Methodology, Software, Validation, Writing – review & editing. RN: Investigation, Validation, Writing – review & editing. JF: Funding acquisition, Project administration. JH: Conceptualization, Funding acquisition, Project administration, Resources, Supervision, Validation, Writing – original draft, Writing – review & editing.

## Funding

This research was funded by the German Research Foundation (DFG) Schwerpunktprogramm (special priority programme) 1158 ‘Antarctic Research with Comparative Investigations in Arctic Ice areas’ (project number 501756173) as well as by the Sonderforschungsbereiche Transregio-Programme 212 (NC3, project numbers 316099922 and 396774617). It was also supported by core funding from the Natural Environment Research Council to the British Antarctic Survey’s Ecosystems Program. Support for the article processing charge was granted by the DFG and the Open Access Publication Fund of Bielefeld University.

## Acknowledgements

We thank Claire Stainfield, Camille Toscani, and Cameron Fox-Clarke for their assistance in the field.

## Conflict of interest

The authors declare that the research was conducted in the absence of any commercial or financial relationships that could be construed as a potential conflict of interest.

## Notes

### Competing Interest Statement

The authors have declared no competing interest.

