## Supplementary for "Skin but not gut microbial communities in Antarctic fur seals vary with social density"

Petroula Botsidou


- Supplementary figures
  - **Figure S1.**
  - **Figure S2.**
  - **Figure S3.**
- Supplementary tables
  - **Table S1.**
  - **Table S2.**
  - **Table S3.**
  - **Table S4.**
  - **Table S5.**
  - **Table S6.**
  - **Table S7.**
  - **Table S8.**
  - **Table S9.**
  - **Table S10.**


### Supplementary figures

#### Figure S1.

Rarefaction curves for skin (A) and gut (B) samples showing the number of observed ASVs in relation to the sequencing depth of every sample.

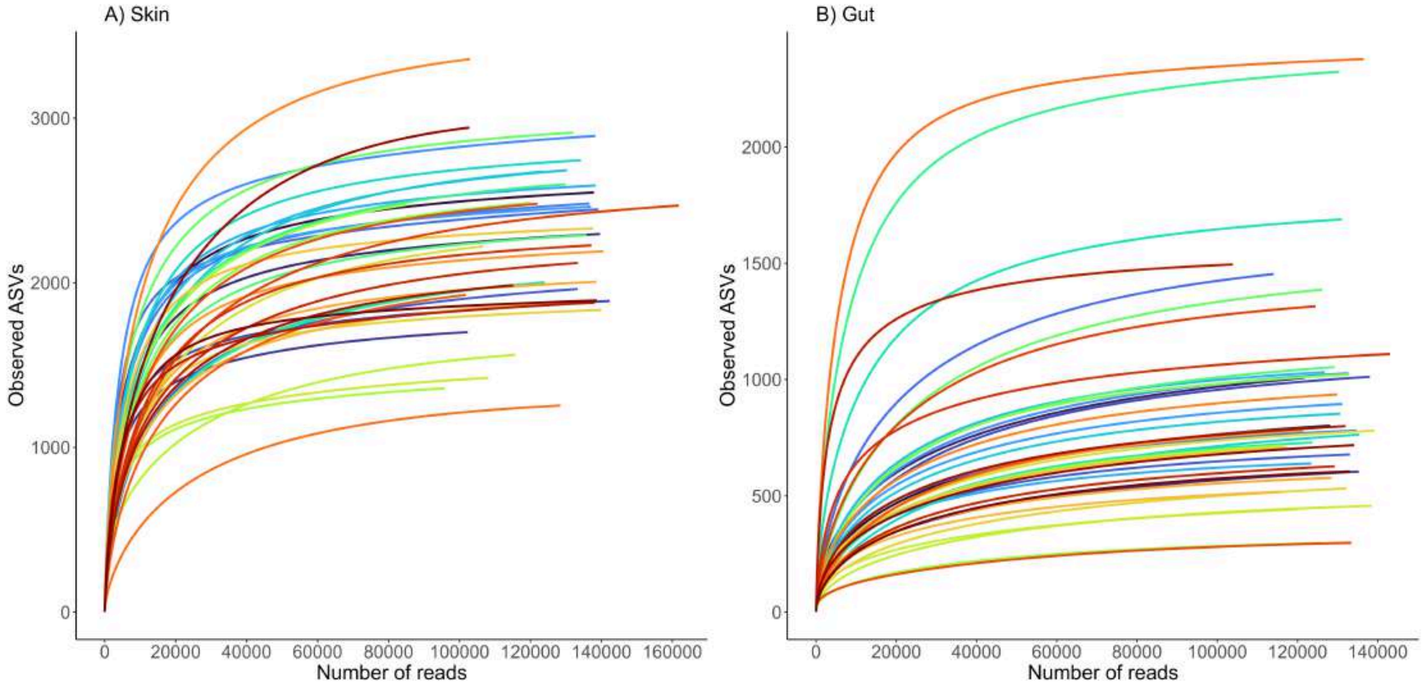

#### Figure S2.

Distribution of Shannon diversity index for skin and gut samples presented with boxplots.

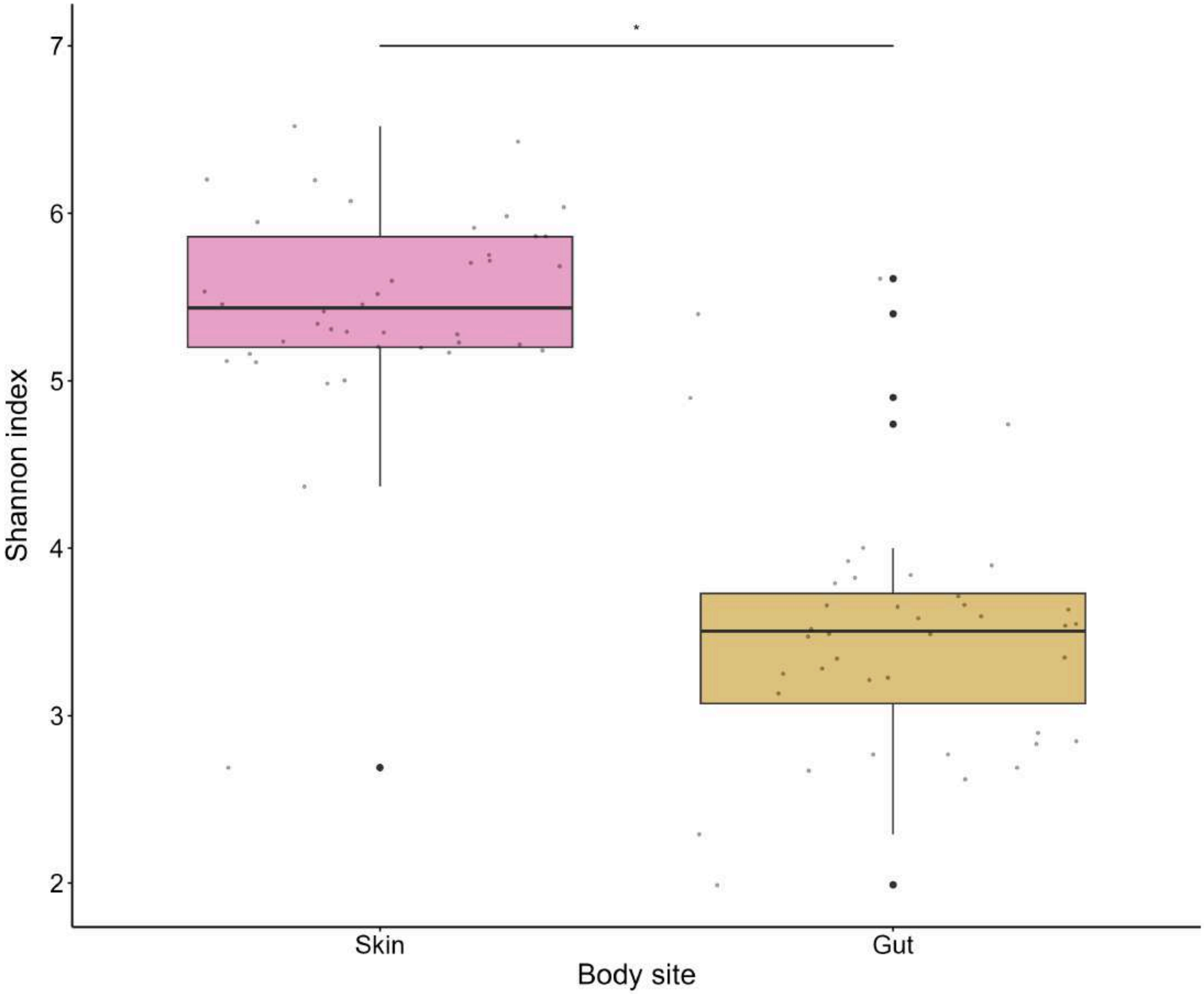

#### Figure S3.

Shannon diversity index for skin samples including outlier (pup H11).

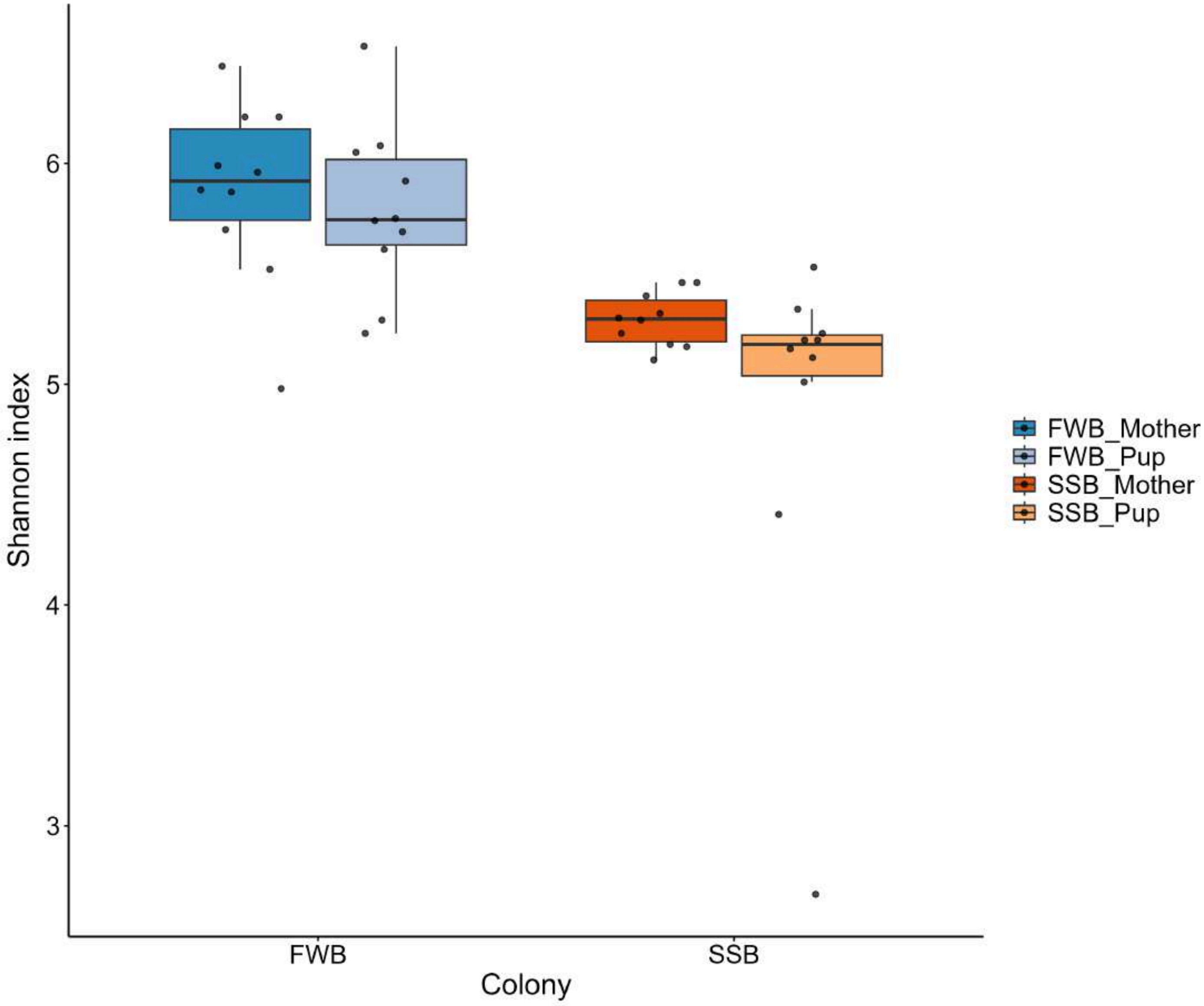


### Supplementary tables

#### Table S1.

Sample IDs together with their corresponding metadata.

| Sample_ID | Body_site | Colony | Age | ID_Pair | Sex |
|---|---|---|---|---|---|
| C1 | Skin | FWB | Pup | F1 | M |
| C1 | Gut | FWB | Pup | F1 | M |
| C17 | Skin | FWB | Pup | F17 | M |
| C17 | Gut | FWB | Pup | F17 | M |
| C18 | Skin | FWB | Pup | F18 | F |
| C18 | Gut | FWB | Pup | F18 | F |
| C21 | Skin | FWB | Pup | F21 | M |
| C21 | Gut | FWB | Pup | F21 | M |

| Sample_ID | Body_site | Colony | Age | ID_Pair | Sex |
|---|---|---|---|---|---|
| C22 | Skin | FWB | Pup | F22 | M |
| C22 | Gut | FWB | Pup | F22 | M |
| C25 | Skin | FWB | Pup | F25 | F |
| C25 | Gut | FWB | Pup | F25 | F |
| C3 | Skin | FWB | Pup | F3 | F |
| C3 | Gut | FWB | Pup | F3 | F |
| C5 | Skin | FWB | Pup | F5 | M |
| C5 | Gut | FWB | Pup | F5 | M |
| C6 | Skin | FWB | Pup | F6 | F |
| C6 | Gut | FWB | Pup | F6 | F |
| C7 | Skin | FWB | Pup | F7 | M |
| C7 | Gut | FWB | Pup | F7 | M |
| F1 | Skin | FWB | Mother | C1 | F |
| F1 | Gut | FWB | Mother | C1 | F |
| F17 | Skin | FWB | Mother | C17 | F |
| F17 | Gut | FWB | Mother | C17 | F |
| F18 | Skin | FWB | Mother | C18 | F |
| F18 | Gut | FWB | Mother | C18 | F |
| F21 | Skin | FWB | Mother | C21 | F |
| F21 | Gut | FWB | Mother | C21 | F |
| F22 | Skin | FWB | Mother | C22 | F |
| F22 | Gut | FWB | Mother | C22 | F |
| F25 | Skin | FWB | Mother | C25 | F |
| F25 | Gut | FWB | Mother | C25 | F |
| F3 | Skin | FWB | Mother | C3 | F |
| F3 | Gut | FWB | Mother | C3 | F |
| F5 | Skin | FWB | Mother | C5 | F |
| F5 | Gut | FWB | Mother | C5 | F |
| F6 | Skin | FWB | Mother | C6 | F |
| F6 | Gut | FWB | Mother | C6 | F |
| F7 | Skin | FWB | Mother | C7 | F |
| F7 | Gut | FWB | Mother | C7 | F |
| H11 | Skin | SSB | Pup | S12 | M |
| H11 | Gut | SSB | Pup | S12 | M |
| H13 | Skin | SSB | Pup | S14 | M |
| H13 | Gut | SSB | Pup | S14 | M |
| H15 | Skin | SSB | Pup | S16 | F |
| H15 | Gut | SSB | Pup | S16 | F |
| H16 | Skin | SSB | Pup | S17 | F |
| H16 | Gut | SSB | Pup | S17 | F |
| H18 | Skin | SSB | Pup | S19 | M |
| H18 | Gut | SSB | Pup | S19 | M |
| H19 | Skin | SSB | Pup | S20 | F |
| H19 | Gut | SSB | Pup | S20 | F |
| H2 | Skin | SSB | Pup | S5 | F |
| H2 | Gut | SSB | Pup | S5 | F |
| H20 | Skin | SSB | Pup | S21 | M |
| H20 | Gut | SSB | Pup | S21 | M |
| H21 | Skin | SSB | Pup | S22 | F |
| H21 | Gut | SSB | Pup | S22 | F |
| H22 | Skin | SSB | Pup | S23 | F |
| H22 | Gut | SSB | Pup | S23 | F |
| S12 | Skin | SSB | Mother | H11 | F |
| S12 | Gut | SSB | Mother | H11 | F |
| S14 | Skin | SSB | Mother | H13 | F |

| Sample_ID | Body_site | Colony | Age | ID_Pair | Sex |
|---|---|---|---|---|---|
| S14 | Gut | SSB | Mother | H13 | F |
| S16 | Skin | SSB | Mother | H15 | F |
| S16 | Gut | SSB | Mother | H15 | F |
| S17 | Skin | SSB | Mother | H16 | F |
| S17 | Gut | SSB | Mother | H16 | F |
| S19 | Skin | SSB | Mother | H18 | F |
| S19 | Gut | SSB | Mother | H18 | F |
| S20 | Skin | SSB | Mother | H19 | F |
| S20 | Gut | SSB | Mother | H19 | F |
| S21 | Skin | SSB | Mother | H20 | F |
| S21 | Gut | SSB | Mother | H20 | F |
| S22 | Skin | SSB | Mother | H21 | F |
| S22 | Gut | SSB | Mother | H21 | F |
| S23 | Skin | SSB | Mother | H22 | F |
| S23 | Gut | SSB | Mother | H22 | F |
| S5 | Skin | SSB | Mother | H2 | F |
| S5 | Gut | SSB | Mother | H2 | F |
| FWB soil | Control | NA | NA | NA | NA |
| SSB soil | Control | NA | NA | NA | NA |
| FWB air | Control | NA | NA | NA | NA |
| SSB air | Control | NA | NA | NA | NA |
| Gloves | Control | NA | NA | NA | NA |
| Hands | Control | NA | NA | NA | NA |

### Table S2.

Summary of total reads and ASVs for each sample before and after filtering steps.

| Sample_ID | Reads_before | ASVs_before | Reads_after | ASVs_after | Colony | Age | Pair_ID | Sex |
|---|---|---|---|---|---|---|---|---|
| **Skin** | | | | | | | | |
| S1 | 137872 | 2548 | 102058 | 1675 | FWB | Mother | P1 | F |
| S2 | 139601 | 2296 | 127035 | 1658 | FWB | Mother | P2 | F |
| S3 | 102354 | 1700 | 84223 | 1187 | FWB | Mother | P3 | F |
| S4 | 142368 | 1889 | 120277 | 1231 | FWB | Mother | P4 | F |
| S5 | 133357 | 1962 | 128985 | 1523 | FWB | Mother | P5 | F |
| S6 | 139142 | 2446 | 106173 | 1566 | FWB | Mother | P6 | F |
| S7 | 136650 | 2481 | 119116 | 1688 | FWB | Mother | P7 | F |
| S8 | 138275 | 2892 | 109980 | 1920 | FWB | Mother | P8 | F |
| S9 | 137085 | 2462 | 121119 | 1761 | FWB | Mother | P9 | F |
| S10 | 138423 | 2591 | 127313 | 1906 | FWB | Mother | P10 | F |
| S11 | 130377 | 2684 | 122930 | 2112 | FWB | Pup | P1 | M |
| S12 | 123451 | 2675 | 116922 | 1964 | FWB | Pup | P2 | M |
| S13 | 134272 | 2744 | 122955 | 1986 | FWB | Pup | P3 | F |
| S14 | 113392 | 1984 | 111418 | 1713 | FWB | Pup | P4 | M |
| S15 | 124015 | 2004 | 122480 | 1772 | FWB | Pup | P5 | M |
| S16 | 129911 | 2598 | 124469 | 2122 | FWB | Pup | P6 | F |
| S17 | 135878 | 2289 | 128200 | 1781 | FWB | Pup | P7 | F |
| S18 | 132064 | 2912 | 112272 | 2161 | FWB | Pup | P8 | M |
| S19 | 119875 | 2484 | 115733 | 2036 | FWB | Pup | P9 | F |
| S20 | 85319 | 2475 | 81363 | 1884 | FWB | Pup | P10 | M |
| S41 | 95783 | 1359 | 79198 | 1022 | SSB | Mother | P11 | F |
| S42 | 115688 | 1562 | 69159 | 1006 | SSB | Mother | P12 | F |
| S43 | 108110 | 1422 | 94681 | 1059 | SSB | Mother | P13 | F |

| Sample_ID | Reads_before | ASVs_before | Reads_after | ASVs_after | Colony | Age | Pair_ID | Sex |
|---|---|---|---|---|---|---|---|---|
| S44 | 106722 | 2221 | 104037 | 1806 | SSB | Mother | P14 | F |
| S45 | 139973 | 1835 | 130412 | 1257 | SSB | Mother | P15 | F |
| S46 | 137719 | 2330 | 118673 | 1478 | SSB | Mother | P16 | F |
| S47 | 132514 | 1883 | 122940 | 1492 | SSB | Mother | P17 | F |
| S48 | 138623 | 2006 | 133725 | 1539 | SSB | Mother | P18 | F |
| S49 | 140594 | 2191 | 129234 | 1530 | SSB | Mother | P19 | F |
| S50 | 102971 | 3358 | 53257 | 1357 | SSB | Mother | P20 | F |
| S51 | 128440 | 1254 | 118420 | 708 | SSB | Pup | P11 | M |
| S52 | 101788 | 1926 | 99039 | 1511 | SSB | Pup | P12 | M |
| S53 | 121989 | 2480 | 117808 | 2011 | SSB | Pup | P13 | F |
| S54 | 161803 | 2469 | 157676 | 1993 | SSB | Pup | P14 | F |
| S55 | 137203 | 2227 | 132252 | 1726 | SSB | Pup | P15 | M |
| S56 | 133413 | 2120 | 130611 | 1764 | SSB | Pup | P16 | F |
| S57 | 138054 | 1879 | 116671 | 1259 | SSB | Pup | P17 | F |
| S58 | 115139 | 1984 | 112402 | 1667 | SSB | Pup | P18 | M |
| S59 | 102756 | 2942 | 96133 | 1791 | SSB | Pup | P19 | F |
| S60 | 138753 | 1892 | 125142 | 1327 | SSB | Pup | P20 | F |
| **Gut** | | | | | | | | |
| S21 | 128133 | 802 | 123735 | 693 | FWB | Mother | P1 | F |
| S22 | 121795 | 1007 | 119253 | 857 | FWB | Mother | P2 | F |
| S23 | 135288 | 604 | 130053 | 501 | FWB | Mother | P3 | F |
| S24 | 138047 | 1012 | 136810 | 812 | FWB | Mother | P4 | F |
| S25 | 133107 | 678 | 128822 | 558 | FWB | Mother | P5 | F |
| S26 | 114083 | 1454 | 111551 | 1097 | FWB | Mother | P6 | F |
| S27 | 134726 | 781 | 131082 | 606 | FWB | Mother | P7 | F |
| S28 | 132666 | 1027 | 131237 | 844 | FWB | Mother | P8 | F |
| S29 | 131133 | 895 | 119005 | 747 | FWB | Mother | P9 | F |
| S30 | 123419 | 639 | 115566 | 549 | FWB | Mother | P10 | F |
| S31 | 126797 | 1032 | 125141 | 798 | FWB | Pup | P1 | M |
| S32 | 130674 | 853 | 128000 | 708 | FWB | Pup | P2 | M |
| S33 | 135348 | 763 | 134367 | 598 | FWB | Pup | P3 | F |
| S34 | 131071 | 1689 | 123160 | 1208 | FWB | Pup | P4 | M |
| S35 | 123697 | 729 | 117175 | 548 | FWB | Pup | P5 | M |
| S36 | 130421 | 2324 | 118851 | 1490 | FWB | Pup | P6 | F |
| S37 | 129274 | 1054 | 127912 | 829 | FWB | Pup | P7 | F |
| S38 | 126113 | 1387 | 120236 | 1063 | FWB | Pup | P8 | M |
| S39 | 132930 | 1022 | 130513 | 736 | FWB | Pup | P9 | F |
| S40 | 125826 | 297 | 123578 | 261 | FWB | Pup | P10 | M |
| S61 | 115927 | 716 | 114144 | 594 | SSB | Mother | P11 | F |
| S62 | 119924 | 442 | 115951 | 384 | SSB | Mother | P12 | F |
| S63 | 138493 | 457 | 138133 | 393 | SSB | Mother | P13 | F |
| S64 | 139148 | 780 | 134705 | 609 | SSB | Mother | P14 | F |
| S65 | 132141 | 532 | 131508 | 451 | SSB | Mother | P15 | F |
| S66 | 117274 | 708 | 115771 | 556 | SSB | Mother | P16 | F |
| S67 | 115728 | 517 | 113255 | 422 | SSB | Mother | P17 | F |
| S68 | 128483 | 577 | 126712 | 497 | SSB | Mother | P18 | F |
| S69 | 121406 | 780 | 118184 | 590 | SSB | Mother | P19 | F |
| S70 | 129922 | 936 | 127375 | 733 | SSB | Mother | P20 | F |
| S71 | 136512 | 2378 | 114932 | 1390 | SSB | Pup | P11 | M |
| S72 | 124953 | 793 | 124043 | 622 | SSB | Pup | P12 | M |
| S73 | 133403 | 298 | 126803 | 218 | SSB | Pup | P13 | F |
| S74 | 124463 | 1315 | 115110 | 848 | SSB | Pup | P14 | F |
| S75 | 143075 | 1110 | 131069 | 666 | SSB | Pup | P15 | M |
| S76 | 129303 | 627 | 125755 | 512 | SSB | Pup | P16 | F |
| S77 | 103914 | 1495 | 68707 | 636 | SSB | Pup | P17 | F |

| Sample_ID | Reads_before | ASVs_before | Reads_after | ASVs_after | Colony | Age | Pair_ID | Sex |
|---|---|---|---|---|---|---|---|---|
| S78 | 132013 | 800 | 126028 | 677 | SSB | Pup | P18 | M |
| S79 | 134140 | 718 | 128959 | 609 | SSB | Pup | P19 | F |
| S80 | 133087 | 605 | 130486 | 520 | SSB | Pup | P20 | F |

### Table S3.

Total reads and ASVs for each body site (skin versus gut) before filtering steps.

| Type | Total_reads | Mean_reads | SD_reads | Min_reads | Max_reads | Total_ASVs | Mean_ASVs | SD_ASVs | Min_ASVs | Max_ASVs |
|---|---|---|---|---|---|---|---|---|---|---|
| Skin & Gut | 10209573 | 127619.7 | 12576.77 | 85319 | 161803 | 33624 | 1576.11 | 805.77 | 297 | 3358 |
| Skin | 5071716 | 126792.9 | 16029.29 | 85319 | 161803 | 27720 | 2236.40 | 460.30 | 1254 | 3358 |
| Gut | 5137857 | 128446.4 | 7878.19 | 103914 | 143075 | 11674 | 915.82 | 457.13 | 297 | 2378 |

### Table S4.

Total reads and ASVs for each body site (skin versus gut) after filtering steps.

| Type | Total_reads | Mean_reads | SD_reads | Min_reads | Max_reads | Total_ASVs | Mean_ASVs | SD_ASVs | Min_ASVs | Max_ASVs |
|---|---|---|---|---|---|---|---|---|---|---|
| Skin & Gut | 9470168 | 118377.1 | 16595.84 | 53257 | 157676 | 7336 | 1154.74 | 563.14 | 218 | 2161 |
| Skin | 4546491 | 113662.3 | 19626.96 | 53257 | 157676 | 6946 | 1623.72 | 345.00 | 708 | 2161 |
| Gut | 4923677 | 123091.9 | 11273.38 | 68707 | 138133 | 3719 | 685.75 | 268.67 | 218 | 1490 |

### Table S5.

Total counts for each phylum for each body site (skin versus gut).

| Phylum | Reads | ASVs | Relative abundance (%) |
|---|---|---|---|
| **Skin** | | | |
| Pseudomonadota | 1368453 | 1988 | 30.13 |
| Bacillota | 1178581 | 1116 | 25.95 |
| Actinomycetota | 790466 | 896 | 17.41 |
| Bacteroidota | 661194 | 2049 | 14.56 |
| Fusobacteriota | 362651 | 53 | 7.99 |
| Patescibacteria | 34873 | 85 | 0.77 |
| Deinococcota | 22556 | 30 | 0.50 |
| Campylobacterota | 21560 | 35 | 0.47 |
| Chloroflexota | 18179 | 85 | 0.40 |
| Acidobacteriota | 17039 | 147 | 0.38 |
| Gemmatimonadota | 12923 | 76 | 0.28 |
| Thermodesulfobacteriota | 12551 | 32 | 0.28 |
| Cyanobacteria | 11903 | 34 | 0.26 |
| Deferribacterota | 7356 | 10 | 0.16 |
| Verrucomicrobiota | 6703 | 74 | 0.15 |
| Planctomycetota | 4379 | 31 | 0.10 |
| Myxococcota | 4131 | 62 | 0.09 |
| Bdellovibrionota | 2221 | 34 | 0.05 |
| Halobacterota | 844 | 3 | 0.02 |
| Abditibacteriota | 481 | 6 | 0.01 |
| Dependentiae | 411 | 2 | 0.01 |
| Nitrospirota | 661 | 9 | 0.01 |

| Phylum | Reads | ASVs | Relative abundance (%) |
|---|---|---|---|
| Spirochaetota | 592 | 6 | 0.01 |
| Synergistota | 425 | 5 | 0.01 |
| Armatimonadota | 14 | 1 | 0.00 |
| Caldisericota | 29 | 1 | 0.00 |
| Cloacimonadota | 38 | 2 | 0.00 |
| Fibrobacterota | 85 | 3 | 0.00 |
| Latescibacterota | 10 | 1 | 0.00 |
| Sumerlaeota | 196 | 2 | 0.00 |
| **Gut** | | | |
| Bacillota | 1783757 | 780 | 36.23 |
| Pseudomonadota | 902317 | 1074 | 18.33 |
| Bacteroidota | 786854 | 1027 | 15.98 |
| Fusobacteriota | 767847 | 38 | 15.59 |
| Actinomycetota | 405457 | 495 | 8.23 |
| Campylobacterota | 251746 | 28 | 5.11 |
| Thermodesulfobacteriota | 6157 | 15 | 0.13 |
| Patescibacteria | 4920 | 54 | 0.10 |
| Deferribacterota | 4311 | 7 | 0.09 |
| Acidobacteriota | 3029 | 61 | 0.06 |
| Deinococcota | 2170 | 18 | 0.04 |
| Cyanobacteria | 1353 | 16 | 0.03 |
| Verrucomicrobiota | 1245 | 19 | 0.03 |
| Gemmatimonadota | 1164 | 23 | 0.02 |
| Chloroflexota | 637 | 30 | 0.01 |
| Abditibacteriota | 52 | 1 | 0.00 |
| Bdellovibrionota | 203 | 5 | 0.00 |
| Cloacimonadota | 6 | 1 | 0.00 |
| Halobacterota | 29 | 1 | 0.00 |
| Myxococcota | 205 | 11 | 0.00 |
| Nitrospirota | 52 | 4 | 0.00 |
| Planctomycetota | 75 | 4 | 0.00 |
| Spirochaetota | 14 | 3 | 0.00 |
| Sumerlaeota | 17 | 2 | 0.00 |
| Synergistota | 60 | 2 | 0.00 |

### Table S6.

Core skin microbiota calculated as the shared ASVs among 90% of the individuals.

| ASV | Phylum | Family | Genus | Species | Prevalence | Reads | Relative abundance (%) |
|---|---|---|---|---|---|---|---|
| ASV_1 | Fusobacteriota | Fusobacteriaceae | Fusobacterium | mortiferum | 40 | 136109 | 5.96 |
| ASV_2 | Fusobacteriota | Fusobacteriaceae | Fusobacterium | perfoetens | 40 | 97930 | 4.29 |
| ASV_3 | Pseudomonadota | Moraxellaceae | Psychrobacter | NA | 40 | 59742 | 2.61 |
| ASV_4 | Bacillota | Clostridiaceae | Clostridium sensu stricto 2 | NA | 40 | 101402 | 4.44 |
| ASV_5 | Pseudomonadota | Enterobacteriaceae | Escherichia-Shigella | coli | 40 | 37876 | 1.66 |
| ASV_6 | Fusobacteriota | Leptotrichiaceae | Oceanivirga | NA | 40 | 5316 | 0.23 |
| ASV_7 | Actinomycetota | Coriobacteriaceae | Collinsella | stercoris | 40 | 77603 | 3.40 |
| ASV_8 | Bacillota | Peptostreptococcaceae | Peptoclostridium | NA | 40 | 69971 | 3.06 |
| ASV_10 | Pseudomonadota | Moraxellaceae | Psychrobacter | NA | 40 | 110080 | 4.82 |

| ASV | Phylum | Family | Genus | Species | Prevalence | Reads | Relative abundance (%) |
|---|---|---|---|---|---|---|---|
| ASV_12 | Bacillota | Clostridiaceae | Clostridium sensu stricto 2 | NA | 40 | 63041 | 2.76 |
| ASV_13 | Pseudomonadota | Moraxellaceae | Psychrobacter | urativorans | 40 | 97401 | 4.26 |
| ASV_14 | Bacillota | Ruminococcaceae | Paludicola | NA | 40 | 49739 | 2.18 |
| ASV_16 | Fusobacteriota | Fusobacteriaceae | Fusobacterium | NA | 40 | 37404 | 1.64 |
| ASV_17 | Bacillota | Butyricicoccaceae | NA | NA | 40 | 51276 | 2.24 |
| ASV_19 | Pseudomonadota | Moraxellaceae | Psychrobacter | arcticus | 40 | 64623 | 2.83 |
| ASV_20 | Bacillota | Lachnospiraceae | [Eubacterium] fissicatena group | NA | 40 | 35946 | 1.57 |
| ASV_27 | Bacillota | Lachnospiraceae | Lachnoclostridium | NA | 40 | 32777 | 1.43 |
| ASV_29 | Actinomycetota | Micrococcaceae | Paeniglutamicibacter | NA | 40 | 55043 | 2.41 |
| ASV_30 | Pseudomonadota | Moraxellaceae | Psychrobacter | NA | 40 | 44080 | 1.93 |
| ASV_33 | Bacteroidota | Bacteroidaceae | Bacteroides | NA | 40 | 19876 | 0.87 |
| ASV_36 | Bacillota | Lachnospiraceae | [Ruminococcus] gnavus group | NA | 40 | 14768 | 0.65 |
| ASV_37 | Bacillota | Lachnospiraceae | Blautia | NA | 40 | 34333 | 1.50 |
| ASV_38 | Pseudomonadota | Moraxellaceae | Psychrobacter | maritimus | 40 | 33904 | 1.48 |
| ASV_40 | Actinomycetota | Intrasporangiaceae | Knoellia | NA | 40 | 42241 | 1.85 |
| ASV_41 | Actinomycetota | Micrococcaceae | Arthrobacter | psychrochitiniphilus | 40 | 39053 | 1.71 |
| ASV_46 | Bacillota | Peptostreptococcaceae | NA | NA | 40 | 28782 | 1.26 |
| ASV_48 | Bacteroidota | Bacteroidaceae | Bacteroides | NA | 40 | 8779 | 0.38 |
| ASV_51 | Bacillota | Lachnospiraceae | Blautia | hansenii | 40 | 16597 | 0.73 |
| ASV_52 | Pseudomonadota | Moraxellaceae | Psychrobacter | cryohalolentis | 40 | 23313 | 1.02 |
| ASV_53 | Bacillota | Family XI | Peptoniphilus | methioninivorax | 40 | 1735 | 0.08 |
| ASV_54 | Bacillota | Clostridiaceae | Clostridium sensu stricto 1 | perfringens | 40 | 22164 | 0.97 |
| ASV_55 | Actinomycetota | Intrasporangiaceae | Intrasporangium | NA | 40 | 30156 | 1.32 |
| ASV_56 | Bacillota | [Eubacterium] coprostanoligenes group | NA | NA | 40 | 11225 | 0.49 |
| ASV_62 | Pseudomonadota | Moraxellaceae | Psychrobacter | cryohalolentis | 40 | 20518 | 0.90 |
| ASV_66 | Bacillota | Clostridiaceae | Clostridium sensu stricto 1 | perfringens | 40 | 13968 | 0.61 |
| ASV_67 | Bacillota | Lachnospiraceae | Tuzzerella | NA | 40 | 9163 | 0.40 |
| ASV_75 | Bacillota | Oscillospiraceae | UCG-005 | NA | 40 | 8609 | 0.38 |
| ASV_76 | Pseudomonadota | Pasteurellaceae | Otariodibacter | NA | 40 | 6068 | 0.27 |
| ASV_94 | Pseudomonadota | Pasteurellaceae | Otariodibacter | NA | 40 | 12951 | 0.57 |
| ASV_101 | Pseudomonadota | Moraxellaceae | Acinetobacter | NA | 40 | 13064 | 0.57 |
| ASV_102 | Bacteroidota | Weeksellaceae | Ornithobacterium | NA | 40 | 12764 | 0.56 |
| ASV_116 | Pseudomonadota | Comamonadaceae | Polaromonas | NA | 40 | 11279 | 0.49 |
| ASV_119 | Deinococcota | Deinococcaceae | Deinococcus | marmoris | 40 | 12058 | 0.53 |
| ASV_121 | Pseudomonadota | Neisseriaceae | Neisseria | zalophi | 40 | 12828 | 0.56 |
| ASV_122 | Campylobacterota | Helicobacteraceae | Helicobacter | typhlonius | 40 | 10983 | 0.48 |
| ASV_141 | Bacillota | Family XI | Anaerococcus | NA | 40 | 10327 | 0.45 |
| ASV_145 | Thermodesulfobacteriota | Desulfovibrionaceae | NA | NA | 40 | 8153 | 0.36 |
| ASV_157 | Pseudomonadota | Moraxellaceae | Psychrobacter | NA | 40 | 9708 | 0.42 |
| ASV_168 | Pseudomonadota | Moraxellaceae | Psychrobacter | NA | 40 | 8166 | 0.36 |
| ASV_170 | Bacillota | Ruminococcaceae | Fournierella | NA | 40 | 3142 | 0.14 |
| ASV_184 | Pseudomonadota | Rhodanobacteraceae | Dokdonella | NA | 40 | 6794 | 0.30 |
| ASV_185 | Deferribacterota | Deferribacteraceae | Mucispirillum | schaedleri | 40 | 7046 | 0.31 |
| ASV_229 | Pseudomonadota | Enterobacteriaceae | Enterobacter | hormaechei | 40 | 3174 | 0.14 |
| ASV_236 | Pseudomonadota | Moraxellaceae | Acinetobacter | baumannii | 40 | 3693 | 0.16 |
| ASV_241 | Actinomycetota | Nocardiaceae | Rhodococcus | yunnanensis | 40 | 5276 | 0.23 |

| ASV | Phylum | Family | Genus | Species | Prevalence | Reads | Relative abundance (%) |
|---|---|---|---|---|---|---|---|
| ASV_242 | Pseudomonadota | Moraxellaceae | Psychrobacter | NA | 40 | 4278 | 0.19 |
| ASV_260 | Bacillota | Family XI | Tissierella | NA | 40 | 4232 | 0.19 |
| ASV_318 | Bacillota | Peptostreptococcaceae | Romboutsia | NA | 40 | 2940 | 0.13 |
| ASV_346 | Pseudomonadota | Rhodanobacteraceae | Rhodanobacter | NA | 40 | 2984 | 0.13 |
| ASV_516 | Pseudomonadota | Alcaligenaceae | Achromobacter | NA | 40 | 1372 | 0.06 |
| ASV_11 | Bacillota | Family XI | Helcococcus | NA | 39 | 4890 | 0.21 |
| ASV_18 | Bacillota | Family XI | Ezakiella | massiliensis | 39 | 3681 | 0.16 |
| ASV_35 | Bacillota | Family XI | Anaerococcus | NA | 39 | 2484 | 0.11 |
| ASV_39 | Fusobacteriota | Fusobacteriaceae | Fusobacterium | mortiferum | 39 | 18143 | 0.79 |
| ASV_43 | Bacteroidota | Bacteroidaceae | Bacteroides | NA | 39 | 8628 | 0.38 |
| ASV_49 | Pseudomonadota | Moraxellaceae | Psychrobacter | NA | 39 | 24184 | 1.06 |
| ASV_58 | Bacillota | Ruminococcaceae | Faecalibacterium | NA | 39 | 3509 | 0.15 |
| ASV_64 | Bacillota | Oscillospiraceae | UCG-002 | NA | 39 | 6037 | 0.26 |
| ASV_80 | Bacteroidota | Bacteroidaceae | Bacteroides | NA | 39 | 4179 | 0.18 |
| ASV_81 | Bacillota | Lachnospiraceae | Tyzzerella | NA | 39 | 6365 | 0.28 |
| ASV_83 | Pseudomonadota | Moraxellaceae | Psychrobacter | cryohalolentis | 39 | 14042 | 0.61 |
| ASV_96 | Bacillota | Clostridiaceae | Clostridium sensu stricto 1 | NA | 39 | 11566 | 0.51 |
| ASV_108 | Bacillota | Oscillospiraceae | UCG-005 | NA | 39 | 3160 | 0.14 |
| ASV_109 | Pseudomonadota | Moraxellaceae | Moraxella | NA | 39 | 12830 | 0.56 |
| ASV_118 | Bacillota | Lactobacillaceae | Levilactobacillus | NA | 39 | 12191 | 0.53 |
| ASV_120 | Actinomycetota | Micrococcaceae | Arthrobacter | agilis | 39 | 11818 | 0.52 |
| ASV_127 | Actinomycetota | Dermacoccaceae | Allobranchiibius | NA | 39 | 11953 | 0.52 |
| ASV_134 | Bacillota | Oscillospiraceae | UCG-005 | NA | 39 | 4873 | 0.21 |
| ASV_147 | Pseudomonadota | Moraxellaceae | Alkanindiges | NA | 39 | 8848 | 0.39 |
| ASV_179 | Bacillota | Acidaminococcaceae | Phascolarctobacterium | NA | 39 | 1926 | 0.08 |
| ASV_182 | Bacillota | Family XI | Parvimonas | NA | 39 | 3341 | 0.15 |
| ASV_218 | Actinomycetota | Nocardioidaceae | Nocardioides | NA | 39 | 6213 | 0.27 |
| ASV_240 | Actinomycetota | Dermacoccaceae | Allobranchiibius | NA | 39 | 5668 | 0.25 |
| ASV_262 | Actinomycetota | Nakamurellaceae | Nakamurella | panacisegetis | 39 | 4656 | 0.20 |
| ASV_265 | Bacillota | Family XI | Finegoldia | magna | 39 | 3701 | 0.16 |
| ASV_274 | Actinomycetota | Micrococcaceae | Arthrobacter | psychrochitiniphilus | 39 | 4360 | 0.19 |
| ASV_289 | Pseudomonadota | Moraxellaceae | Psychrobacter | NA | 39 | 4273 | 0.19 |
| ASV_310 | Actinomycetota | Propionibacteriaceae | Tessaracoccus | NA | 39 | 3576 | 0.16 |
| ASV_331 | Bacteroidota | Weeksellaceae | Ornithobacterium | NA | 39 | 2713 | 0.12 |
| ASV_23 | Bacillota | Aerococcaceae | Atopobacter | phocae | 38 | 6841 | 0.30 |
| ASV_28 | Bacteroidota | Bacteroidaceae | Bacteroides | NA | 38 | 19297 | 0.84 |
| ASV_32 | Pseudomonadota | Moraxellaceae | Psychrobacter | urativorans | 38 | 49008 | 2.15 |
| ASV_44 | Bacillota | Family XI | W5053 | NA | 38 | 1376 | 0.06 |
| ASV_45 | Pseudomonadota | Moraxellaceae | Psychrobacter | NA | 38 | 38417 | 1.68 |
| ASV_144 | Bacillota | Oscillospiraceae | NK4A214 group | NA | 38 | 1682 | 0.07 |
| ASV_166 | Actinomycetota | Intrasporangiaceae | Knoellia | NA | 38 | 8947 | 0.39 |
| ASV_176 | Actinomycetota | Intrasporangiaceae | NA | NA | 38 | 8194 | 0.36 |
| ASV_200 | Actinomycetota | Kineosporiaceae | Quadrisphaera | NA | 38 | 6665 | 0.29 |
| ASV_211 | Bacillota | Clostridiaceae | Clostridium sensu stricto 1 | paraputrificum | 38 | 4963 | 0.22 |
| ASV_215 | Bacteroidota | Flavobacteriaceae | Flavobacterium | antarcticum | 38 | 5407 | 0.24 |
| ASV_228 | Pseudomonadota | Moraxellaceae | Psychrobacter | NA | 38 | 4337 | 0.19 |
| ASV_231 | Actinomycetota | Nocardiaceae | Nocardia | NA | 38 | 5674 | 0.25 |
| ASV_251 | Bacillota | Ruminococcaceae | Fournierella | NA | 38 | 935 | 0.04 |
| ASV_252 | Bacillota | Streptococcaceae | Streptococcus | marimammalium | 38 | 4511 | 0.20 |
| ASV_322 | Pseudomonadota | Moraxellaceae | Psychrobacter | NA | 38 | 2332 | 0.10 |
| ASV_350 | Actinomycetota | Dermacoccaceae | Allobranchiibius | NA | 38 | 3236 | 0.14 |

| ASV | Phylum | Family | Genus | Species | Prevalence | Reads | Relative abundance (%) |
|---|---|---|---|---|---|---|---|
| ASV_413 | Actinomycetota | Corynebacteriaceae | Corynebacterium | phocae | 38 | 2567 | 0.11 |
| ASV_444 | Bacillota | Lachnospiraceae | NA | NA | 38 | 1769 | 0.08 |
| ASV_456 | Pseudomonadota | Enterobacteriaceae | Klebsiella | pneumoniae | 38 | 1221 | 0.05 |
| ASV_468 | Bacillota | Ruminococcaceae | Faecalibacterium | prausnitzii | 38 | 1434 | 0.06 |
| ASV_485 | Bacillota | Peptostreptococcaceae | Romboutsia | ilealis | 38 | 946 | 0.04 |
| ASV_665 | Actinomycetota | Nocardiaceae | Nocardia | NA | 38 | 1212 | 0.05 |
| ASV_853 | Actinomycetota | Intrasporangiaceae | NA | NA | 38 | 1016 | 0.04 |
| ASV_1621 | Pseudomonadota | Moraxellaceae | Psychrobacter | NA | 38 | 424 | 0.02 |
| ASV_24 | Actinomycetota | Actinomycetaceae | Arcanobacterium | NA | 37 | 4006 | 0.18 |
| ASV_60 | Bacillota | Aerococcaceae | Atopobacter | phocae | 37 | 3162 | 0.14 |
| ASV_72 | Bacillota | Family XI | Peptoniphilus | NA | 37 | 1131 | 0.05 |
| ASV_78 | Bacteroidota | Rikenellaceae | Alistipes | NA | 37 | 4214 | 0.18 |
| ASV_79 | Bacillota | Planococcaceae | Sporosarcina | NA | 37 | 19551 | 0.86 |
| ASV_126 | Actinomycetota | Micrococcaceae | Arthrobacter | alpinus | 37 | 11735 | 0.51 |
| ASV_150 | Bacillota | Oscillospiraceae | UCG-005 | NA | 37 | 2408 | 0.11 |
| ASV_165 | Pseudomonadota | Pasteurellaceae | Otariodibacter | NA | 37 | 8238 | 0.36 |
| ASV_201 | Bacillota | Peptostreptococcaceae | Terrisporobacter | NA | 37 | 4906 | 0.21 |
| ASV_203 | Actinomycetota | Ilumatobacteraceae | Ilumatobacter | NA | 37 | 6639 | 0.29 |
| ASV_226 | Actinomycetota | Geodermatophilaceae | Antricoccus | NA | 37 | 5521 | 0.24 |
| ASV_230 | Bacillota | Ruminococcaceae | Faecalibacterium | prausnitzii | 37 | 3522 | 0.15 |
| ASV_268 | Pseudomonadota | Moraxellaceae | Psychrobacter | NA | 37 | 3892 | 0.17 |
| ASV_303 | Actinomycetota | Dermacoccaceae | Allobranchiibius | NA | 37 | 4065 | 0.18 |
| ASV_304 | Cyanobacteria | Leptolyngbyaceae | NA | NA | 37 | 3304 | 0.14 |
| ASV_323 | Actinomycetota | Demequinaceae | NA | NA | 37 | 2798 | 0.12 |
| ASV_438 | Bacillota | Lachnospiraceae | Lachnospiraceae NK4A136 group | NA | 37 | 2125 | 0.09 |
| ASV_477 | Actinomycetota | Iamiaceae | Iamia | NA | 37 | 2194 | 0.10 |
| ASV_713 | Bacillota | Ruminococcaceae | Fournierella | NA | 37 | 1170 | 0.05 |
| ASV_140 | Bacillota | Streptococcaceae | Streptococcus | marimammalium | 37 | 9303 | 0.41 |
| ASV_244 | Bacteroidota | Flavobacteriaceae | Flavobacterium | phocarum | 37 | 4787 | 0.21 |

#### Table S7.

Core gut microbiota calculated as the shared ASVs among 90% of the individuals.

| ASV | Phylum | Family | Genus | Species | Prevalence | Reads | Relative abundance (%) |
|---|---|---|---|---|---|---|---|
| ASV_1 | Fusobacteriota | Fusobacteriaceae | Fusobacterium | mortiferum | 40 | 224354 | 8.24 |
| ASV_2 | Fusobacteriota | Fusobacteriaceae | Fusobacterium | perfoetens | 40 | 216034 | 7.93 |
| ASV_3 | Pseudomonadota | Moraxellaceae | Psychrobacter | NA | 40 | 219772 | 8.07 |
| ASV_4 | Bacillota | Clostridiaceae | Clostridium sensu stricto 2 | NA | 40 | 125286 | 4.60 |
| ASV_5 | Pseudomonadota | Enterobacteriaceae | Escherichia-Shigella | coli | 40 | 137668 | 5.06 |
| ASV_6 | Fusobacteriota | Leptotrichiaceae | Oceanivirga | NA | 40 | 163092 | 5.99 |
| ASV_7 | Actinomycetota | Coriobacteriaceae | Collinsella | stercoris | 40 | 86743 | 3.19 |
| ASV_8 | Bacillota | Peptostreptococcaceae | Peptoclostridium | NA | 40 | 83974 | 3.08 |
| ASV_10 | Pseudomonadota | Moraxellaceae | Psychrobacter | NA | 40 | 23945 | 0.88 |
| ASV_11 | Bacillota | Family XI | Helcococcus | NA | 40 | 136068 | 5.00 |
| ASV_12 | Bacillota | Clostridiaceae | Clostridium sensu stricto 2 | NA | 40 | 71694 | 2.63 |

| ASV | Phylum | Family | Genus | Species | Prevalence | Reads | Relative abundance (%) |
|---|---|---|---|---|---|---|---|
| ASV_16 | Fusobacteriota | Fusobacteriaceae | Fusobacterium | NA | 40 | 70609 | 2.59 |
| ASV_17 | Bacillota | Butyricicoccaceae | NA | NA | 40 | 48359 | 1.78 |
| ASV_18 | Bacillota | Family XI | Ezakiella | massiliensis | 40 | 88551 | 3.25 |
| ASV_20 | Bacillota | Lachnospiraceae | [Eubacterium] fissicatena group | NA | 40 | 47149 | 1.73 |
| ASV_27 | Bacillota | Lachnospiraceae | Lachnoclostridium | NA | 40 | 29117 | 1.07 |
| ASV_30 | Pseudomonadota | Moraxellaceae | Psychrobacter | NA | 40 | 11663 | 0.43 |
| ASV_37 | Bacillota | Lachnospiraceae | Blautia | NA | 40 | 17343 | 0.64 |
| ASV_38 | Pseudomonadota | Moraxellaceae | Psychrobacter | maritimus | 40 | 14037 | 0.52 |
| ASV_39 | Fusobacteriota | Fusobacteriaceae | Fusobacterium | mortiferum | 40 | 29788 | 1.09 |
| ASV_41 | Actinomycetota | Micrococcaceae | Arthrobacter | psychrochitiniphilus | 40 | 5289 | 0.19 |
| ASV_51 | Bacillota | Lachnospiraceae | Blautia | hansenii | 40 | 17484 | 0.64 |
| ASV_53 | Bacillota | Family XI | Peptoniphilus | methioninivorax | 40 | 32052 | 1.18 |
| ASV_56 | Bacillota | [Eubacterium] coprostanoligenes group | NA | NA | 40 | 20122 | 0.74 |
| ASV_66 | Bacillota | Clostridiaceae | Clostridium sensu stricto 1 | perfringens | 40 | 10487 | 0.39 |
| ASV_67 | Bacillota | Lachnospiraceae | Tuzzerella | NA | 40 | 16479 | 0.61 |
| ASV_75 | Bacillota | Oscillospiraceae | UCG-005 | NA | 40 | 12914 | 0.47 |
| ASV_229 | Pseudomonadota | Enterobacteriaceae | Enterobacter | hormaechei | 40 | 2189 | 0.08 |
| ASV_236 | Pseudomonadota | Moraxellaceae | Acinetobacter | baumannii | 40 | 1965 | 0.07 |
| ASV_14 | Bacillota | Ruminococcaceae | Paludicola | NA | 39 | 82335 | 3.02 |
| ASV_19 | Pseudomonadota | Moraxellaceae | Psychrobacter | arcticus | 39 | 16096 | 0.59 |
| ASV_29 | Actinomycetota | Micrococcaceae | Paeniglutamicibacter | NA | 39 | 6161 | 0.23 |
| ASV_33 | Bacteroidota | Bacteroidaceae | Bacteroides | NA | 39 | 38046 | 1.40 |
| ASV_35 | Bacillota | Family XI | Anaerococcus | NA | 39 | 53953 | 1.98 |
| ASV_36 | Bacillota | Lachnospiraceae | [Ruminococcus] gnavus group | NA | 39 | 36488 | 1.34 |
| ASV_44 | Bacillota | Family XI | W5053 | NA | 39 | 38582 | 1.42 |
| ASV_46 | Bacillota | Peptostreptococcaceae | NA | NA | 39 | 10312 | 0.38 |
| ASV_54 | Bacillota | Clostridiaceae | Clostridium sensu stricto 1 | perfringens | 39 | 9359 | 0.34 |
| ASV_70 | Bacillota | Aerococcaceae | Atopobacter | phocae | 39 | 21907 | 0.80 |
| ASV_72 | Bacillota | Family XI | Peptoniphilus | NA | 39 | 22606 | 0.83 |
| ASV_13 | Pseudomonadota | Moraxellaceae | Psychrobacter | urativorans | 38 | 23753 | 0.87 |
| ASV_40 | Actinomycetota | Intrasporangiaceae | Knoellia | NA | 38 | 3466 | 0.13 |
| ASV_49 | Pseudomonadota | Moraxellaceae | Psychrobacter | NA | 38 | 6574 | 0.24 |
| ASV_58 | Bacillota | Ruminococcaceae | Faecalibacterium | NA | 38 | 27403 | 1.01 |
| ASV_62 | Pseudomonadota | Moraxellaceae | Psychrobacter | cryohalolentis | 38 | 4897 | 0.18 |
| ASV_64 | Bacillota | Oscillospiraceae | UCG-002 | NA | 38 | 22208 | 0.82 |
| ASV_68 | Bacillota | Family XI | Peptoniphilus | NA | 38 | 24112 | 0.89 |
| ASV_81 | Bacillota | Lachnospiraceae | Tyzzerella | NA | 38 | 13712 | 0.50 |
| ASV_108 | Bacillota | Oscillospiraceae | UCG-005 | NA | 38 | 12993 | 0.48 |
| ASV_116 | Pseudomonadota | Comamonadaceae | Polaromonas | NA | 38 | 2003 | 0.07 |
| ASV_516 | Pseudomonadota | Alcaligenaceae | Achromobacter | NA | 38 | 676 | 0.02 |
| ASV_22 | Campylobacterota | Campylobacteraceae | Campylobacter | blaseri | 37 | 72903 | 2.68 |
| ASV_24 | Actinomycetota | Actinomycetaceae | Arcanobacterium | NA | 37 | 65696 | 2.41 |
| ASV_28 | Bacteroidota | Bacteroidaceae | Bacteroides | NA | 37 | 42842 | 1.57 |
| ASV_43 | Bacteroidota | Bacteroidaceae | Bacteroides | NA | 37 | 31897 | 1.17 |
| ASV_61 | Bacteroidota | Porphyromonadaceae | Porphyromonas | NA | 37 | 29438 | 1.08 |
| ASV_115 | Actinomycetota | Actinomycetaceae | Actinomyces | neuii | 37 | 13729 | 0.50 |
| ASV_122 | Campylobacterota | Helicobacteraceae | Helicobacter | typhlonius | 37 | 2294 | 0.08 |
| ASV_124 | Bacillota | Family XI | NA | NA | 37 | 11796 | 0.43 |

| ASV | Phylum | Family | Genus | Species | Prevalence | Reads | Relative abundance (%) |
|---|---|---|---|---|---|---|---|
| ASV_170 | Bacillota | Ruminococcaceae | Fournierella | NA | 37 | 5907 | 0.22 |
| ASV_185 | Deferribacterota | Deferribacteraceae | Mucispirillum | schaedleri | 37 | 1280 | 0.05 |
| ASV_242 | Pseudomonadota | Moraxellaceae | Psychrobacter | NA | 37 | 1405 | 0.05 |
| ASV_318 | Bacillota | Peptostreptococcaceae | Romboutsia | NA | 37 | 899 | 0.03 |
| ASV_337 | Pseudomonadota | Moraxellaceae | Enhydrobacter | NA | 37 | 596 | 0.02 |
| ASV_823 | Bacillota | Clostridiaceae | Clostridium sensu stricto 1 | NA | 37 | 558 | 0.02 |

### Table S8.

Output of the linear mixed effect models.

| Shannon index | Estimate | SE | Df | CI_lower | CI_upper | Chisq | Pr(>F) |
|---|---|---|---|---|---|---|---|
| **Skin** | | | | | | | |
| Intercept | 5.95 | 0.14 | 28.31 | 5.67 | 6.26 | - | - |
| Colony (SSB) | -0.74 | 0.18 | 1 | -1.11 | -0.38 | 17.51 | <0.001 |
| Age (Pup) | -0.24 | 0.14 | 1 | -0.52 | 0.03 | 3.08 | 0.08 |
| **Gut** | | | | | | | |
| Intercept | 3.52 | 0.21 | 32.5 | 3.11 | 3.95 | - | - |
| Colony (SSB) | -0.07 | 0.25 | 1 | -0.59 | 0.39 | 0.09 | 0.761 |
| Age (Pup) | 0.06 | 0.24 | 1 | -0.41 | 0.57 | 0.07 | 0.792 |

### Table S9.

Output of the linear models.

| Shannon index | Estimate | SE | Df | CI_lower | CI_upper | F-value | P-value |
|---|---|---|---|---|---|---|---|
| **Skin-pups** | | | | | | | |
| Intercept | 6.09 | 0.25 | 1 | 5.56 | 6.63 | - | - |
| Colony (SSB) | -1 | 0.28 | 1 | -1.58 | -0.42 | 11.06 | <0.001 |
| Sex (Male) | -0.51 | 0.28 | 1 | -1.09 | 0.08 | 3.36 | 0.08 |
| **Gut-pups** | | | | | | | |
| Intercept | 3.52 | 0.44 | 1 | 2.59 | 4.46 | - | - |
| Colony (SSB) | 0.07 | 0.48 | 1 | -0.95 | 1.09 | 0.02 | 0.883 |
| Sex (Male) | -0.02 | 0.48 | 1 | -1.04 | 1.01 | 1e-03 | 0.975 |

### Table S10.

Output of PERMANOVA analysis and dispersion test.

| | Permanova | | | | | betadisper | |
|---|---|---|---|---|---|---|---|
| Parameter | Df | SumOfSqs | R2 | F (P) | Pr(>F) (P) | F (b) | Pr(>F) (b) |
| **Skin** | | | | | | | |
| Colony | 1 | 1.662 | 0.228 | 15.462 | 0.001 | 1.832 | 0.184 |
| Age | 1 | 0.292 | 0.04 | 2.716 | 0.009 | 11.132 | 0.002 |
| Pair_ID | 18 | 3.277 | 0.451 | 1.694 | 0.001 | - | - |
| Residual | 19 | 2.042 | 0.281 | - | - | - | - |
| Total | 39 | 7.272 | 1 | - | - | - | - |

| | Permanova | | | | | betadisper | |
|---|---|---|---|---|---|---|---|
| Parameter | Df | SumOfSqs | R2 | F (P) | Pr(>F) (P) | F (b) | Pr(>F) (b) |
| **Gut** | | | | | | | |
| Colony | 1 | 0.601 | 0.081 | 3.829 | 0.001 | 0.334 | 0.567 |
| Age | 1 | 0.695 | 0.094 | 4.431 | 0.001 | 18.564 | <0.001 |
| Pair_ID | 18 | 3.118 | 0.422 | 1.104 | 0.113 | - | - |
| Residual | 19 | 2.982 | 0.403 | - | - | - | - |
| Total | 39 | 7.396 | 1 | - | - | - | - |
| **Skin-pups** | | | | | | | |
| Sex | 1 | 0.139 | 0.046 | 1.12 | 0.243 | 2.129 | 0.162 |
| Colony | 1 | 0.803 | 0.263 | 6.466 | 0.001 | 0.033 | 0.858 |
| Residual | 17 | 2.11 | 0.691 | - | - | - | - |
| Total | 19 | 3.052 | 1 | - | - | - | - |
| **Gut-pups** | | | | | | | |
| Sex | 1 | 0.208 | 0.055 | 1.117 | 0.268 | 0.207 | 0.654 |
| Colony | 1 | 0.439 | 0.115 | 2.354 | 0.003 | 0.002 | 0.963 |
| Residual | 17 | 3.172 | 0.83 | - | - | - | - |
| Total | 19 | 3.82 | 1 | - | - | - | - |
